# The efficacy of different preprocessing steps in reducing motion-related confounds in diffusion MRI connectomics

**DOI:** 10.1101/2020.03.25.008979

**Authors:** Stuart Oldham, Aurina Arnatkevic̆iūtė, Robert E. Smith, Jeggan Tiego, Mark A. Bellgrove, Alex Fornito

## Abstract

Head motion is a major confounding factor in neuroimaging studies. While numerous studies have investigated how motion impacts estimates of functional connectivity, the effects of motion on structural connectivity measured using diffusion MRI have not received the same level of attention, despite the fact that, like functional MRI, diffusion MRI relies on elaborate preprocessing pipelines that require multiple choices at each step. Here, we report a comprehensive analysis of how these choices influence motion-related contamination of structural connectivity estimates. Using a healthy adult sample (*N* = 252), we evaluated 240 different preprocessing pipelines, devised using plausible combinations of different choices related to explicit head motion correction, tractography propagation algorithms, track seeding methods, track termination constraints, quantitative metrics derived for each connectome edge, and parcellations. We found that an approach to motion correction that includes outlier replacement and within-slice volume correction led to a dramatic reduction in cross-subject correlations between head motion and structural connectivity strength, and that motion contamination is more severe when quantifying connectivity strength using mean tract fractional anisotropy rather than streamline count. We also show that the choice of preprocessing strategy can significantly influence subsequent inferences about network organization, with the location of network hubs varying considerably depending on the specific preprocessing steps applied. Our findings indicate that the impact of motion on structural connectivity can be successfully mitigated using recent motion-correction algorithms that include outlier replacement and within-slice motion correction.

**Highlights:**

- We assess how motion affects structural connectivity in 240 preprocessing pipelines
- Motion contamination of structural connectivity depends on preprocessing choices
- Advanced motion correction tools reduce motion confounds
- FA edge weighting is more susceptible to motion effects than streamline count

---

Generating comprehensive maps of brain connectivity, called connectomes (Sporns, Tononi, & Kötter, 2005), has become one of the central goals of neuroscience. In humans, diffusion magnetic resonance imaging (dMRI) is the most widely used technique for studying the anatomical connectivity of the brain. Typically, the outcomes of a tractography experiment are combined with a brain parcellation to generate a whole-brain connectivity matrix, which can then be analysed using the tools of graph theory (Bullmore & Sporns, 2009; Fornito, Zalesky, & Bullmore, 2016). This approach has generated new insights into principles of brain network organization (Bullmore & Sporns, 2009, 2012; van den Heuvel & Sporns, 2013), development (Bassett, Xia, & Satterthwaite, 2018; Cao, Huang, & He, 2017; Morgan, White, Bullmore, & Vértes, 2018; Oldham & Fornito, 2019; Zhao, Xu, & He, 2019), how network anatomy constrains function (Goñi et al., 2014; Honey, Thivierge, & Sporns, 2010; Skudlarski et al., 2008), and how brain connectivity is affected by disease (Crossley et al., 2014; Fornito, Zalesky, & Breakspear, 2015; Stam, 2014).

A major problem for any *in vivo* brain imaging experiment is participant head motion. The potential artefacts caused by motion have been studied extensively for functional MRI, where numerous motion estimation & correction strategies have been proposed (Ciric et al., 2017; Fair et al., 2013; Parkes, Fulcher, Yücel, & Fornito, 2018; Power, Barnes, Snyder, Schlaggar, & Petersen, 2012; Satterthwaite et al., 2013, 2012). It has been widely acknowledged that motion can induce numerous artefacts in dMRI––including misalignment between slice acquisitions, attenuation of signal intensities, and signal dropout––that conspire to limit the accuracy of diffusion signal models and subsequent tractography (Anderson & Gore, 1994; Jones & Basser, 2004; Le Bihan, Poupon, Amadon, & Lethimonnier, 2006). However, the consequences of head motion for dMRI estimates of connectivity, and the efficacy of methods to address them, have not received the same scrutiny as in the functional MRI literature.

Studies examining motion-related contamination of dMRI measures have mostly considered effects on voxel-wise estimates of popular metrics, such as fractional anisotropy (FA). A common finding is that motion can inflate the FA of low anisotropy regions, while diminishing the FA of high anisotropy regions (Jones & Basser, 2004; Le Bihan et al., 2006; Tijssen, Jansen, & Backes, 2009). Residual motion effects can still contaminate data even when motion correction techniques, such as realignment of diffusion volumes, are employed (Ling et al., 2012; Liu, Zhu, & Zhong, 2015; Oguz et al., 2014; Yendiki, Koldewyn, Kakunoori, Kanwisher, & Fischl, 2014). One recent study (Baum et al., 2018) examined how motion affects dMRI measures of pairwise structural connectivity between regions, as typically analysed in a connectomic study. The authors found that motion reduced the strength of connections that are consistently found across people, while increasing the strengths of connections that are inconsistent and/or short-range. They also found that the effects of motion were contingent on the specific tractography algorithm employed, such that motion exerted a larger effect on connectivity estimates of inconsistently detected edges for connectomes generated using a deterministic tractography algorithm compared to probabilistic tractography. This result suggests that the type of tractography algorithm used will influence the extent to which motion biases estimates of structural connectivity.

Choice of tractography algorithm is not the only decision to make in a dMRI connectomic analysis. Indeed, such analyses rely on elaborate, multi-stage preprocessing pipelines that depend on decisions made at each stage. It is plausible that such choices may also influence the degree to which head motion contaminates the resulting connectivity estimates. For instance, one step with an obvious impact is the rigid realignment of each diffusion volume to a common reference (usually the first *b* = 0, or an average of all *b* = 0 images; a step sometimes referred to as motion correction) (Andersson & Skare, 2002; Rohde, Barnett, Basser, Marenco, & Pierpaoli, 2004). Different algorithms are available for performing this step, each varying in their sophistication and efficacy in dealing with other problems, such as signal outliers and within-volume head motion (Andersson et al., 2017; Andersson, Graham, Zsoldos, & Sotiropoulos, 2016; Andersson & Sotiropoulos, 2016).

Other preprocessing steps in dMRI pipelines are not designed to explicitly address head motion confounds but may still influence the severity of motion-related contamination. For example, some techniques filter reconstructed streamlines according to their biological plausibility (e.g., Girard, Whittingstall, Deriche, & Descoteaux, 2014; R. E. Smith, Tournier, Calamante, & Connelly, 2012), such that streamlines do not terminate in inappropriate anatomical regions. Other approaches either filter or reweight streamlines by altering connectivity estimates to more closely match the underlying diffusion signal (Daducci, Dal Palù, Lemkaddem, & Thiran, 2015; Pestilli, Yeatman, Rokem, Kay, & Wandell, 2014; R. E. Smith, Tournier, Calamante, & Connelly, 2013, 2015a). To the extent that motion results in spurious streamlines, such filtering or reweighting methods may be effective in mitigating motion-related confounds in dMRI connectomics, despite not being explicitly intended to do so. Given the wide variety of preprocessing techniques available in dMRI pipelines, a comprehensive analysis of how different preprocessing steps, either alone or in combination, mitigate or exacerbate motion contamination in structural connectivity analyses is required.

In this study, we evaluate the degree to which 16 distinct preprocessing options, and the 240 unique (and sensible) combinations thereof, were successful in mitigating head-motion confounds in dMRI connectomes reconstructed for a sample of 252 individuals. We hypothesized *a priori* that more advanced image realignment methods, such as those that also correct for within-volume motion, combined with methods designed to derive more biologically plausible estimates of structural connectivity (e.g., R. E. Smith et al., 2012, 2015a), would be more effective in mitigating motion-related confounds. Across pipelines, we quantified residual correlations between inter-subject variability in connectivity estimates and motion measures using both the raw structural connectome matrix data and derivative network measures, with the aim of seeing whether particular preprocessing steps were associated with weaker or stronger susceptibility to motion related effects.

## Methods

### Overview

We identified seven key steps in dMRI connectomic pipelines that require investigators to make choices that might affect final estimates of structural connectivity. Each of these steps had 2-3 available choices. A schematic overview of these steps and choices is provided in Figure 1. We constructed distinct preprocessing pipelines based on every feasible combination of choices, resulting in a total of 240 different pipelines. We used this comprehensive, combinatorial approach because specific choices at one stage may interact with choices at other stages. The steps that we focused on are featured prominently in preprocessing pipelines that are implemented as part of the FSL (Jenkinson, Beckmann, Behrens, Woolrich, & Smith, 2012) and MRtrix3 (Tournier et al., 2019) software packages. From FSL, we rely on the preprocessing tools *topup* and *eddy* (Jenkinson et al., 2012), which provide estimation of B0 field inhomogeneities (Andersson, Skare, & Ashburner, 2003; S. M. Smith et al., 2004), and correction for these distortions in addition to correction for motion, eddy current distortions, and signal dropout (Andersson et al., 2017, 2016; Andersson & Sotiropoulos, 2016), respectively. From MRtrix3, we relied on the standard workflow for connectome construction (Tournier, Calamante, & Connelly, 2012; Tournier et al., 2019), due to the availability of many different relevant preprocessing algorithms and choices thereof, particularly those designed to optimize connectomic measures. We focus on these steps and packages because they are commonly used in connectome construction and include many relevant preprocessing choices in a simple workflow. We note that other software packages are available (e.g., DTIstudio, ExploreDTI), and these may entail other preprocessing choices which we do not examine here. Data processing took was conducted using the Multi-modal Australian ScienceS Imaging and Visualisation Environment (MASSIVE) high performance computing infrastructure (Goscinski et al., 2014). Details of each of the steps and choices we consider in our analysis, and the data used, are described in the following sections.

**Figure 1.**
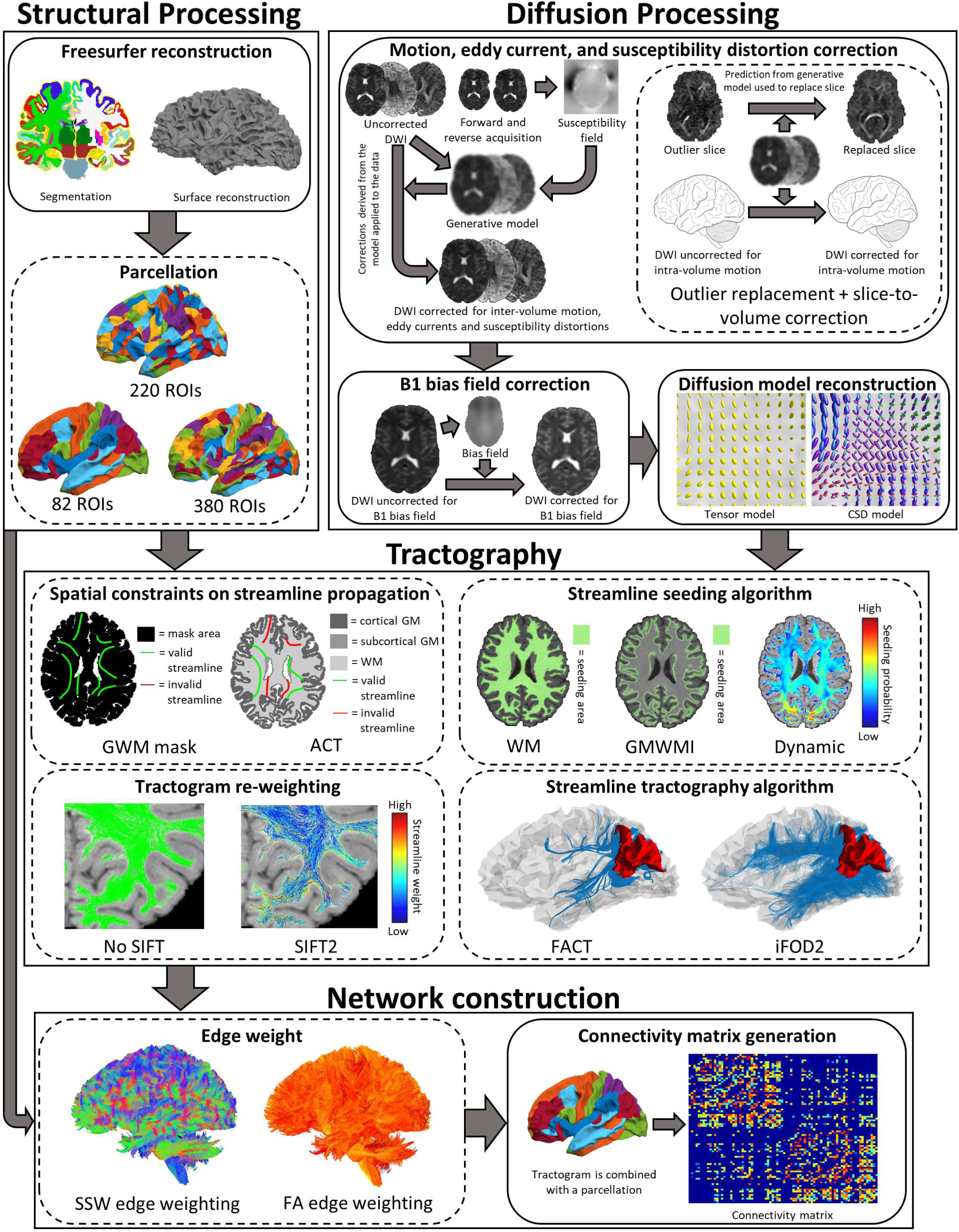
Workflow of preprocessing steps/phases used in structural connectome construction. Square boxes group related steps, round boxes with a continuous border indicate a step used across all pipelines, and round boxes with a dashed border indicate a step requiring choices between different options. Arrows indicate which outputs of preprocessing steps/phases feed into other steps (note in the case of the “Tractography” phase, all the preprocessing steps apart from tractogram filtering occur concurrently, filtering is applied after these steps). Terminology: Constrained spherical deconvolution (CSD), Fibre Assignment by Continuous Tractography (FACT), second-order Integration over Fibre Orientation Distributions (iFOD2), Grey and white matter (GWM), Anatomically Constrained Tractography (ACT), white-matter (WM), grey-matter white-matter interface (GMWMI), Spherical-deconvolution Informed Filtering of Tractograms (SIFT2), sum of streamline weights (SSW), fractional anisotropy (FA). See text for a comprehensive description of each step.

### Participants

A total of 294 healthy participants were recruited as part of a larger study being conducted at Monash University (the participants selected all had both structural and diffusion MRI scans available). Participants were all right-handed, of European ancestry, and had no personal history of neurological or psychiatric disorders, had never suffered loss of consciousness or memory due to head injury, and did not have a history of drug abuse (for further details, see Sabaroedin et al., 2019). The study was conducted in accordance with the Monash University Human Research Ethics Committee (reference number 2012001562). Participants with very large head motion were excluded (further details below), resulting in a final sample of 252 individuals (mean age 23.05 ± 5.28 years, 139 females).

### Image acquisition

Structural and diffusion MRI data were acquired using a Siemens Skyra 3T scanner with a 32-channel head coil at Monash Biomedical Imaging in Clayton, Victoria, Australia. Diffusion data used an interleaved acquisition with the following parameters: 2.5mm^3^ voxel size, TR = 8800ms, TE = 110ms, FOV 240mm, 60 directions with *b* = 3000 s/mm^2^, and seven *b* = 0 s/mm^2^ volumes. In addition, a single *b* = 0 s/mm^2^ was obtained with reversed phase encoding direction for susceptibility field estimation. T1-weighted (T1w) structural scans were acquired using: 1mm^3^ isotropic voxels, TR = 2300ms, TE = 2.07ms, TI = 900ms, and a FOV of 256mm.

### Preprocessing steps common to all pipelines

#### T1w image preprocessing

T1w scans were processed with Freesurfer version 5.3 (Fischl, 2012) to extract cortical surface models of the grey/white matter surface and grey/CSF (pial) surface. All Freesurfer output was checked and, if required, manually corrected for errors in surface reconstruction. The cortical surface models were parcellated to define network nodes for connectomic analysis (parcellations are described in detail below). All images derived from the structural MRI data (including masks and parcellations) were then co-registered to the processed diffusion data, as described below.

#### DWI preprocessing

Diffusion data were processed using MRtrix3 (Tournier et al., 2019) and FSL (Jenkinson et al., 2012).The raw diffusion data were first processed with FSL’s *topup*, using the forward and reverse phase-encoded *b* = 0 s/mm^2^ images toestimate the susceptibility induced off-resonance field (Andersson et al., 2003; S. M. Smith et al., 2004). After motion and eddy current correction, the diffusion data was corrected for B1 field inhomogeneities using FAST in FSL (S. M. Smith et al., 2004; Zhang, Brady, & Smith, 2001).

#### Preprocessing steps varying across pipelines

In this section, we outline the specific choices made at each step of the connectome reconstruction pipeline.

#### Motion and eddy current correction

We compare the effects of two different motion correction strategies implemented by FSL’s *eddy* tool:

1. The first strategy corrected for eddy-induced current distortions, susceptibility-induced distortions (using the field inhomogeneity estimate from *topup*), and inter-volume head motion (Andersson & Sotiropoulos, 2016). Here, a Gaussian Process-based generative model is used to obtain a prediction of each diffusion volume. Comparison of the predicted and actual images yields estimates of eddy current-induced distortions, which are then used to realign the data (Andersson & Sotiropoulos, 2016). This approach is henceforth referred to as EDDY1.
2. The second strategy additionally corrected for outliers in the diffusion signal and within-volume motion (Andersson et al., 2017, 2016). Outliers are detected by comparing the diffision signal to the prediction obtained from the generative model. Any slices where the image intensity is at least four standard deviations lower than the prediction are classified as containing signal dropout, and intensities within that slice are replaced by those produced by the generative model. Within-volume motion is estimated and corrected by incorporating data regarding the timing of acquisition of each image slice and a smooth model of subject rigid-body motion during the acquisition of each image volume into the same generative model. A forward model then uses these estimates of slice-wise movement to reconstruct the corrected diffusion data. Herceforth we refer to this procedure as EDDY2.

#### Streamline tractography algorithm

Diffusion MRI can be used to reconstruct the brain’s white matter pathways using one of several different streamlines tractography algorithms (Jeurissen, Descoteaux, Mori, & Leemans, 2019). While many details of such algorithms can vary substantially, they are commonly assigned to one of two classifications: (1) *Determinstic*: At each streamline vertex, the orientation in which to propagate the streamline is chosen non-stochastically; (2) *Probabilistic*: At each streamline vertex, one of a distribution of plausible orientations is chosen in which to propagate the streamline.

In general, probabilistic algorithms are more sensitive in tracking non-dominant fibre pathways and are prone to false positives; determinsitic algorithms, on the other hand, are more conservative and prone to false neagtives (Reveley et al., 2015; Thomas et al., 2014). This trade-off between specificity and sensitivity can affect network analyses (Zalesky et al., 2016). We thus chose exemplars of each algorithm class to evaluate their relative relationships to motion.

Deterministic tractography was performed using the Fibre Assignment by Continuous Tractography (FACT) algorithm (Mori, Crain, Chacko, & van Zijl, 1999; Mori & van Zijl, 2002) as implaimented in MRtrix3 (Tournier et al., 2019). The algorithm propagates streamlines in the direction of the most collinear fibre orientation estimated within the voxel in which the streamline vertex resides. We definied one fibre orientation in each voxel by estimating the diffusion tensor using iteratively reweighted linear least squares (Veraart, Sijbers, Sunaert, Leemans, & Jeurissen, 2013) and then calculating the primary eigenvector.

Probabilistic tractography was performed using the second-order Integration over Fibre Orientation Distributions (iFOD2) algorithm, as impliamented in MRtrix3 (Tournier et al., 2019), which utilises Fibre Orientation Distributions (FODs) estimated for each voxel using Constrained Spherical Deconvolution (Tournier, Calamante, & Connelly, 2010; Tournier, Calamante, & Connelly, 2007; Tournier et al., 2012). For candidate streamline trajectories emanating from the current vertex, probabilities are calculated based on the amplitudes of the FODs along those trajectories; trajectories with greater probabilities are then more likely to be randomly chosen for streamline propagation. This approach can improve the reconstruction of tracts in highly curved and crossing fibre regions (Tournier et al., 2010, 2012).

For both tractography algorithms, we used a maximum streamline length of 400mm, maximum curvature of 45° per step, and generated 1,000,000 streamlines. Default parameters for each algorithm were used for step size (0.25mm for iFOD2; 1.25mm for FACT) and streamline termination criteria (0.05 FOD amplitude for iFOD2; 0.05 amplitude of the primary eigenvector for FACT).

#### Spatial constraints on streamline propogation

Some spatial constraints are often applied to tractography algorithms to limit the reconstruction of biologically implausible streamlines. A simple constraint is the use of a tissue mask that ensures streamlines are only generated within that mask. One masking stratgey that we evalulated is the use of a binary mask of grey and white matter (GWM) areas, which ensured that streamlines began, traversed, and terminated, within the brain parenchyma. The GWM masks were generated by combining the Freesurfer segmenation of the grey and white matter into a single binary mask.

A second set of constraints we evaluated were implemented as part of the Anatomically Constrained Tractography (ACT) framework. In this approach, the brain is segmented (Patenaude, Smith, Kennedy, & Jenkinson, 2011; S. M. Smith, 2002; S. M. Smith et al., 2004; Zhang et al., 2001) into different tissue types (cortical grey matter, subcortical grey matter, white matter, and cerebrospinal fluid [CSF]). ACT exploits the information provided by the segmentation to ensure that streamlines only trace paths through, and terminate at, anatomically correct tissue locations (e.g. streamlines cannot terminate in white matter regions, or pass through CSF). This approach can reduce false positives in tract reconstruction (R. E. Smith et al., 2012). When combined specifically with the probabilistic iFOD2 algorithm, we utilised the “backtracking” capability of ACT, in which streamlines that are improperly terminated are truncated and a new trajectory is sought (R. E. Smith et al., 2012).We hypothesized that ACT, by removing biologically implausible streamlines, may indirectly mitigates motion confounds.

#### Streamline seeding algorithm

We evaluated three different strategies for seeding streamlines:

1. randomly seeding from voxels in the white matter (WM);
2. randomly seeding streamlines from the grey matter - white matter interface (GMWMI), which has been shown to improve the reconstruction of shorter pathways (R. E. Smith et al., 2013; R. E. Smith, Tournier, Calamante, & Connelly, 2015b); and
3. dynamic seeding, whereby streamlines are seeded preferentially from voxels where the streamline density is under-estimated with respect to fibre density estimates from the diffusion model, helping to improve the reconstruction of those pathways that are more difficult to track (R. E. Smith et al., 2015a).

#### Streamline re-weighting

Commonly, every streamline produced by tractography is interpreted as having the same “weight” as every other streamline. However, the density of streamlines produced during tractography is not necessarily reflective of the density of the underlying white matter fibres, which limits biological interpretability of tractogram-based connectivity estimates. A class of semi-global tractogram re-weighting algorithms seek to modulate the contributions of different streamlines toward the model by weighting individual streamlines in order to minimise this discrepancy (Sherbondy 2008; Sherbondy 2009; Smith 2013; Smith 2015; Daducci 2014; Pestilli 2014). By improving the biological plausibility of connectivity estimates, it is plausible that such approaches may potentially reduce motion-related confounds. Here, we evaluated the effects of both using and not using the second Spherical-deconvolution Informed Filtering of Tractograms (SIFT2) method (R. E. Smith et al., 2015a). SIFT2 operates by modelling the expected fibre densities at every voxel and then weighting each streamline according to how well it fits the model.

#### Parcellation

Construction of a structural brain network necessitates division of the brain grey matter into different regions to represent the distinct nodes of the network. As the choice of parcellation can affect numerous properties of a brain network (Fornito, Zalesky, & Bullmore, 2010; Zalesky et al., 2010), we investigated three different parcellations:

1. *82 nodes*: 34 cortical (Desikan et al., 2006) and seven subcortical regions (Fischl et al., 2002) per hemisphere delineated using sulcal, gyral and other anatomical boundaries.
2. *220 nodes*: random division of each hemisphere into 100 approximately equal sized cortical regions (Fornito et al., 2011; Zalesky 2010), combined with a subcortical parcellation of three striatal (Tziortzi et al., 2014) and seven thalamic (Behrens, Johansen-Berg, et al., 2003; Behrens, Woolrich, et al., 2003) regions. Subcortical regions were non-linearily registered using FSLs *FNIRT* (Jenkinson et al., 2012) to the subject’s T1w image data.
3. *380 nodes*: a recent 180-region cortical parcellation generated by combining information from multiple imaging modalities (Glasser et al., 2016) was combined with the same subcortical regions as the 220 node parcellation to produce a final parcellation of 380 regions.

Cortical parcellations were generated on the Freesurfer-estimated surface models and projected out to the T1w image grid. This volume-based cortical parcellation was then merged with the subcortical parcellations. Following rigid-body coregistration of the subject’s diffusion image to their T1w image, using FSLs *FLIRT* (Greve & Fischl, 2009; Jenkinson, Bannister, Brady, & Smith, 2002; Jenkinson & Smith, 2001), and subsequent inversion of the resulting transformation, each combined parcellation image was mapped to the subject’s diffusion image data in native space.

#### Network construction and edge weight

Connectivity matrices were generated from the tractogram and parcellation data by assigning streamlines to each of the closest regions within a 5 mm radius of the streamline endpoints (R. E. Smith et al., 2015b), yielding an undirected *N* × *N* connectivity matrix, where *N* is the number of regions defined in the parcellation. In this matrix, each element [*i, j*] indicates the presence and some measure of strength of connectivity (i.e. edge weight) between regions *i* and *j*.

Various metrics can be derived from the reconstruction and/or image data to quantify the magnitude of connectivity between each pair of regions (i.e. the weight of the edge). Here we assess two options:

1. Sum of streamline weights (SSW). The weights of each streamline connecting a pair of regions are summed to give the weight of that edge. Note that in the absence of use of the SIFT2 method, this is equivalent to streamline count (i.e. a weight of 1.0 is assumed for all streamlines).
2. Weighting edges by FA values (Baker et al., 2015; Baum et al., 2017; van den Heuvel & Sporns, 2011). Weighting edges in this manner is thought to provide an index of the microstructural integrity of the underlying white matter connections, although this interpretation is likely simplistic (Beaulieu, 2002; Jones, 2010; Jones, Knösche, & Turner, 2013). To obtain these edge weights, streamlines are assigned the mean FA of the voxels they traverse. For streamlines connecting regions/nodes *i* and *j*, the mean of these FA streamline weights is taken (note that when SIFT2 is used, the streamline weights it provides are also incorporated into this computation).

Prior studies suggest that the relationship between motion and edge consistency is stronger in FA rather than SSW edge weighted networks (Baum et al., 2018), and other DWI studies have shown that FA is more susceptible to the effects of motion (Jones & Basser, 2004; Le Bihan et al., 2006; Tijssen et al., 2009).

#### Combining preprocessing choices into distinct pipelines

In summary, the possibilities for combining components into preprocessing pipelines are as follows:

- *Motion correction (MotionCorr)*: EDDY1, EDDY2;
- *Streamline tractography algorithm*: FACT, iFOD2;
- *Spatial constrainsts on streamline propogation*: GWM, ACT;
- *Streamline seeding algorithm (SeedAlgor)*: WM, GMWMI, dynamic;
- *Tractogram re-weighting* : SIFT2, none;
- *Parcellation*: 82, 220 and 380 nodes;
- *Edge weight (EdgeWeight)*: SSW, FA.

We examined every possible combination of these preprocessing steps, which resulted in 240 distinct pipelines (note that because the implementation of GMWMI seeding in *MRtrix3* is dependent upon ACT, pipelines combining GWM and GMWMI were excluded).

#### Quantifying in-scanner head motion

Previous studies investigating the influence of head motion on dMRI measures have quantified motion by measuring the spatial displacement that occurs between volumes acquired through time (Baum et al., 2018; Roalf et al., 2016). A comparable estimate of in-scanner head motion can be directly obtained from *eddy* outputs (Andersson & Sotiropoulos, 2016; Bastiani et al., 2019) by taking the square root of the mean voxelwise squared displacement of each volume. This displacement is calculated for each volume with respect to both the previous volume, and to the first volume; from these, we take the mean across all volumes to obtain scalar estimates of relative (*REL*_*all*_) and absolute (*ABS*_*all*_) head motion. For each pipeline, the corresponding motion parameter estimates from EDDY1 or EDDY2 were utilized. In this article we focus on results obtained using *ABS*_*all*_; the results for *REL*_*all*_, along with five other measures that have also been used to characterise motion (Baum et al., 2018; Roalf et al., 2016), are presented in the supplementary material (Table S1). Note these other measures are calculated based on the transformation matrix used when performing an affine registration on the raw dMRI images.

Six measures of head motion (*ABS*_*all*_ and *REL*_*all*_ for both EDDY1 and EDDY2, and mean absolute/relative *b* = 0 volume-to-volume displacement; see Table S1) were used to identify subjects with excessive head motion. Our criterion was that participants must not exceed 1.5 inter-quartile ranges above the third quartile for *any* measure. We adopted this stringent criterion to focus our experiment on the effects of normal levels of in-scanner head motion. This resulted in removal of 42 participants.

#### Quantifying motion-related contamination of connectivity strength

To quantify the influence of motion on anatomical connectivity, we computed, independently for each edge, a Spearman correlation between edge weight and each specific measure of head motion across participants (Baum et al., 2018). We term this measure the quality control – structural connectivity (QC-SC) correlation. Note that such correlations were computed at an edge consistency-based threshold of 5% – only edges that had a nonzero edge weight in over 5% of participants were analysed – and the correlations were estimated using only those subjects with nonzero connectivity in that specific edge (i.e. zero edge weights were excluded). We evaluated pipeline performance as the proportion of edges in the network for which where there was a significant QC-SC relationship (*p* < .05 uncorrected), as per past work (Parkes et al., 2018). We also examined how QC-SC related to edge length (the mean length of the streamlines connecting two regions), edge *consistency* (the proportion of participants in which a given edge was present), edge *weight variability* (the coefficient of variation in edge weights across participants), and edge length (Baum et al., 2018).

In addition, we considered the impact of thresholding on QC-SC estimates. Thresholding is often implemented when analysing human brain structural networks to remove spurious edges (de Reus & van den Heuvel, 2013; Roberts, Perry, Roberts, Mitchell, & Breakspear, 2017; Rubinov & Sporns, 2010; Zalesky et al., 2016). A common procedure is to exclude edges based on the extent to which reconstruction of that edge varies across individuals. We examined two such approaches:

1. An edge *consistency*-based threshold: considering edges that are non-zero in over some fraction of participants (de Reus 2013), where we considered thresholds varying from 5% to 90%;
2. An edge *variability*-based threshold: considering those edges with the smallest coefficient of variation (CoV) (Roberts et al 2017), where we considered thresholds varying from the 10^th^ to 100^th^ percentiles.

Many connectomic analyses consider properties of the network that are calculated for each *node*, rather than, or in addition to, each edge (Bullmore & Sporns, 2009; Fornito et al., 2016). These properties are also likely to be affected by motion, because they are derived directly from the edge weights. One of the most widely examined properties is nodal centrality - the capacity of a node to influence or be influenced by other nodes – which is used to identify network ‘hubs’ (Fornito et al., 2016; van den Heuvel & Sporns, 2013). Although there are many ways to determine nodal centrality (Oldham et al., 2019), we focus here on node strength (the sum of edge weights assigned to a node) as it is the simplest and most commonly used centrality measure, and it has previously been found to be affected by motion (Baum et al., 2018). To assess how node strength is related to motion we calculated, at each threshold (as defined above), the proportion of nodes whose strength demonstrated a significant correlation with motion (Spearman correlation, *p* < .05 uncorrected). Previous studies have found that the choice of preprocessing steps can result in different network topologies (Li, Rilling, Preuss, Glasser, & Hu, 2012; Yeh, Smith, Dhollander, Calamante, & Connelly, 2019), including the locations of hubs. To this end, we calculated, independently for each pipeline, the mean rank of node strength for each node across all participants; we then quantified, for every possible pair of pipelines, the correlation coefficient between these values across all nodes; we refer to these as “Node strength rank (NSR) correlations”.

## Results

### Basic connectome properties

Across the various reconstruction pipelines, the mean edge density (i.e. fraction of non-zero edges) across participants ranged from 3-94%. As would be expected, the sparsest networks resulted from the combination of FACT and high-resolution parcellations (e.g., the 380-node parcellation), whereas the densest networks resulted from the combination of iFOD2 and low-resolution parcellations (82 nodes).

Figure 2 displays how the basic network properties of edge density (fraction of non-zero edges) and mean edge weight vary across pipelines, when specifically using the 220 node parcellation (data for other parcellations are shown in Figures S1-S2), averaged across individuals (using a 5% edge consistency based threshold). As expected, the edge densities and mean edge weights of connectomes generated with iFOD2 (average density of 74%) were higher than those generated with FACT (average density of 9%). When using both iFOD2 and SSW edge weighting, variations in mean edge weight across pipeline choices were smaller than other combinations, but nonetheless spanned a range of 49 to 78. In contrast, when using FACT, mean edge weight showed greater variability as a function of other pipeline choices. For instance, the combination of ACT with no streamline filtering resulted in a mean edge weight that was approximately double that obtained when employing SIFT2, or when using GWM rather than ACT. By forbidding streamlines from terminating in the white matter, ACT ensures that a greater fraction of reconstructed streamlines contribute to the connectome, thereby increasing the mean edge weight.

**Figure 2.**
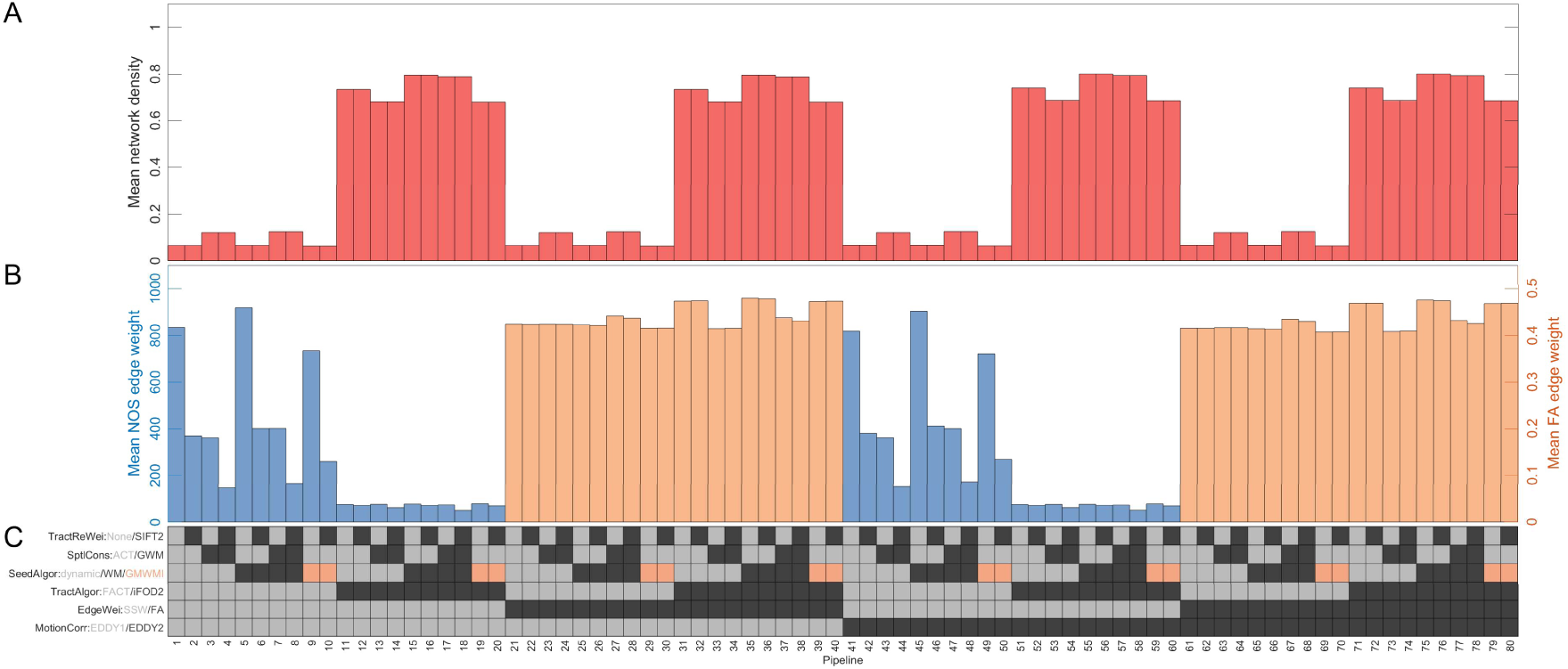
Network density and mean edge-weight for the 220 node parcellation across 80 combinations of preprocessing choices. (**A**) The mean participant network density (y-axis) in each pipeline (x-axis). (**B**) The mean edge weight for SSW (left y-axis) and FA (right y-axis) in each pipeline (x-axis). (**C**) The preprocessing options used in each pipeline. Each row corresponds to a preprocessing step, with the possible options for that step colour-coded; the colour of the squares in each column indicate the specific methods used for a given pipeline.

### Relationship between in-scanner head motion and structural connectivity

Sample estimates of mean head motion are presented in Table S2, and are comparable to those reported in previous analyses (Baum et al., 2018; Roalf et al., 2016). The distribution of scores on each of the motion measures for the sample are displayed in Figure S3. To quantify the effects of motion on structural connectivity, we correlated edge weight and head motion (*ABS*_*all*_) across participants independently at each edge, to obtain an edgewise QC-SC correlation estimate. We then calculated the proportion of edges that had a significant QC-SC correlation at *p* < .05, uncorrected. This proportion varied widely across pipelines, ranging from 2.3% to 83.5%.

Figure 3A shows the proportion of tested edges (i.e. edges that were nonzero in at least 5% of participants) with a significant QC-SC relationship across those pipelines utilising the 220 node parcellation (4.79-71.78%), along with a key to indicate the specific preprocessing steps applied in each case (Figure 3C). This figure suggests that in most cases, there were more significant negative than positive QC-SC correlations; i.e. where estimated motion leads to statistically significant changes in connectivity estimates, greater motion leads to reduced reconstructed connectivity.

**Figure 3.**
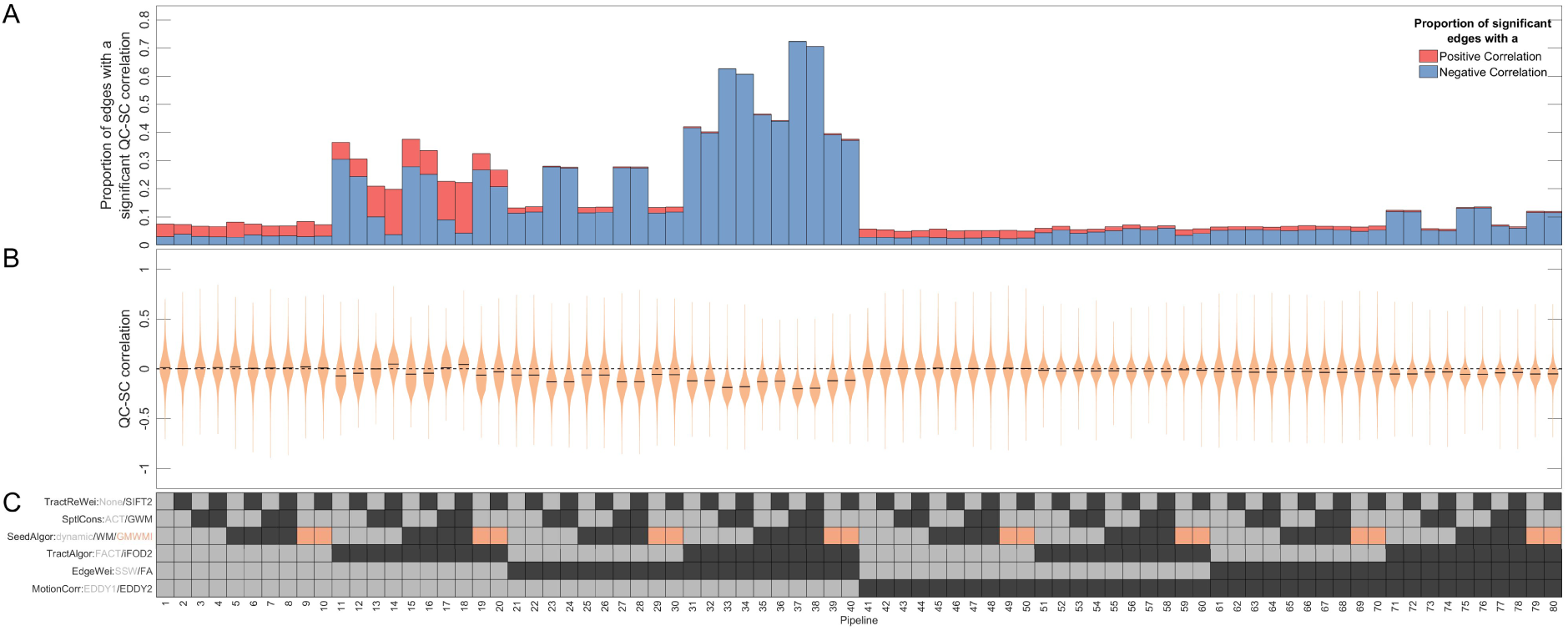
The effect of in-scanner motion (*ABS*_*all*_) on structural connectivity when using a 220 node parcellation. (**A**) The proportion of edges (y-axis) that had a significant (*p* < .05, uncorrected) QC-SC correlation in each pipeline (x-axis). Each bar is coloured to show out of those significant edges, what proportion were negative (blue) or positive (red). (**B**) The full distributions of QC-SC correlations (y-axis) for each pipeline (x-axis). (**C**) The preprocessing options used in each pipeline. Each row corresponds to a preprocessing step, with the possible options for that step colour-coded; the colour of the squares in each column indicate the specific methods used for a given pipeline.

Figure 3B shows the distributions of QC-SC correlations across pipelines. In most cases, the distributions are centred on zero or a negative value, suggesting that increased head motion most often correlates with lower estimated SC. Pipelines exhibiting a negative mode generally used a combination of EDDY1 and SIFT2. Qualitatively similar results were obtained across all parcellations, although connectomes constructed at coarser resolution parcellations suffer less motion-related contamination (Fig S4-S5).

The biggest effect on QC-SC correlations was the choice of motion correction strategy, with EDDY2 dramatically reducing the proportion of significant QC-SC correlations relative to EDDY1, regardless of other preprocessing choices. Compared to EDDY1, EDDY2 comprises two additional capabilities: outlier replacement and within-volume motion correction (Andersson et al., 2017, 2016; Andersson & Sotiropoulos, 2016). To determine which of these was responsible for the reduction in QC-SC correlations, we also evaluated performance for a selected number of pipelines in which *eddy* had been run with outlier replacement only, which we term EDDY1.5. The proportion of significant QC-SC correlations was intermediate between the results of EDDY1 and EDDY2, suggesting that these two capabilities contribute approximately equally toward reducing correlations between connectivity and motion (Figure S6). The difference between EDDY1.5 and EDDY2 was particularly notable for pipelines using iFOD2 and FA-weighting, where EDDY2 reduced the proportion of significant QC-SC correlations from approximately 30% to 12%.

The differences between EDDY1 and EDDY2 were most pronounced when either iFOD2 or FA edge weighting were used. The combination of EDDY1, iFOD2 and FA weighting results in particularly poor performance, with 60-72% of edges showing significant QC-SC correlations. Applying EDDY2 prior to iFOD2 and FA edge weighting reduced the QC-SC proportions to levels comparable to other choices (except when ACT was also used, in which case the proportions were slightly elevated).

The method of constraining streamline propagation also impacted QC-SC correlations. For pipelines that combined EDDY1, iFOD2, and SSW edge weighting, use of ACT displayed greater correlation with motion than otherwise equivalent pipelines that used a simple GWM mask. Conversely, when connectomes were weighted using FA rather than SSW, ACT was associated with a lower proportion of significant QC-SC correlations relative to GWM mask. This trend was reversed again with the use of EDDY2 in conjunction with FA weighting, where ACT increased the proportion of QC-SC correlations relative to the GWM mask. These findings suggest complex interactions are possible between different choices in DWI preprocessing pipelines.

As all of our correlations were conducted using only those subjects with a non-zero connectivity value for each specific edge, the number of data points available for any given QC-SC correlation varied across edges. To check whether omission of subjects with absent connectivity affected our findings, we computed correlations across all subjects for all edges in which at least one subject had registered a connectivity value. The results of this analysis were similar to our main findings; the key exceptions were a weaker average magnitude of QC-SC values, and no differentiation between ACT and the GWM mask when iFOD2, EDDY1 and FA edge weighting were used (Fig S7). In a second analysis, we applied a range of different edge thresholds (5-90% edge consistency and 10^th^-100^th^ percentile edge variability) and calculated the proportion of remaining edges significantly correlated with motion estimates; this analysis revealed a similar pattern of results (Fig S8), suggesting that our primary observations are not due to connectome thresholding effects.

Finally, we also examined QC-SC correlations when alternative measures of head motion were used. As with our main findings, pipelines with FA edge weighting, iFOD2 and EDDY1 displayed notably higher levels of correlation with motion compared to other pipelines (Figures S9-S14). However, when estimating motion with RELb3000 (Figure S13) or TSNR (Figure S14), differences between pipelines using EDDY1 and EDDY2 were much smaller.

#### Relationship between QC-SC correlations and edge metrics

Next, we evaluated how QC-SC correlations relate to edge consistency, edge variability, and edge length. Figure 4 shows these associations for 10 example pipelines (selected as exemplars of the patterns observed across all data). Across all pipelines, short-range edges were most likely to be reconstructed consistently across subjects (Fig 4A-E). The dependence of QC-SC correlations on edge consistency was similar across pipelines, having a funnel-like shape characterised by a wider range of QC-SC correlations for less consistent edges and a smaller range of low-value QC-SC correlations for more consistent edges; in other words, edges that are found more consistently across individuals show a lower range of motion susceptibility.

**Figure 4.**
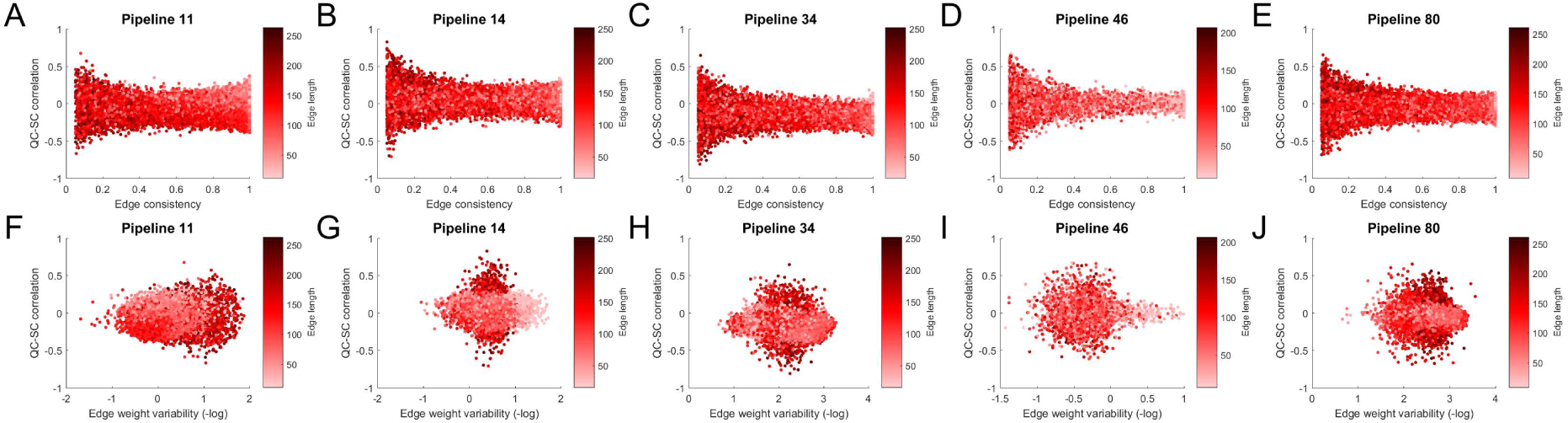
Relationship between QC-SC correlations and edge length, edge consistency, and edge weight variability. The first row shows the relationship between QC-SC correlations (y-axis), edge consistency (x-axis), and edge length (colourmap), while the second shows the relationship between QC-SC correlations(x-axis), edge weight variability (y-axis), and edge length (colourmap). Pipeline 11: EDDY1, SSW edge weighting, iFOD2, dynamic seeding, ACT, no SIFT2. Pipeline 14: EDDY1, SSW edge weighting, iFOD2, dynamic seeding, GWM, SIFT2. Pipeline 34: EDDY1, FA edge weighting, iFOD2, dynamic seeding, GWM, SIFT2. Pipeline 46: EDDY2, SSW edge weighting, iFOD2, WM seeding, ACT, SIFT2. Pipeline 80: EDDY2, FA edge weighting, iFOD2, GMWMI seeding, GWM, SIFT2.

The relationship between QC-SC correlations, edge length, and edge *weight variability* (rather than edge *consistency*) was more complex, with a stronger dependency on pipeline (Fig 4F-J). Pipelines that used iFOD2 and SSW edge weighting showed a wide range of QC-SC correlations for edges with higher weight variability (e.g. pipeline 11; Fig 4F), in line with the edge consistency findings. However, the combination of these choices (i.e., iFOD2 and SSW edge weighting) with SIFT2 led to a larger range of QC-SC correlations for edges with moderate weight variability (e.g. pipelines 14 and 34; Fig 4G and 4H). Pipelines using FACT and SSW edge weighting showed a wide range of QC-SC correlations for edges with lower weight variability (e.g. pipeline 46; Fig 4I). Weighting edges by FA increased the variability of QC-SC correlations for moderate weight variability (e.g. pipeline 80; Fig 4J). Finally, we compared QC-SC correlations to edge weighting, finding that for SSW the *weakest* edges displayed the strongest QC-SC correlations, while for FA the *strongest* edges displayed the strongest QC-SC correlations (Figure S15).

We re-evaluated these relationships after estimating QC-SC correlations across *all* subjects for all edges in which at least one subject had registered a connectivity value (i.e. retaining null connectivity values). The relationship between edge consistency and QC-SC correlation no longer displayed the same funnel shape; instead, this funnel shape was reversed (the most consistent edges displayed the strongest QC-SC relationships) or the relationship was now clearly negative (Figure S16). Additionally, the relationship between edge weight variability and QC-SC correlations showed greater heterogeneity across pipelines.

#### Relationship between head motion and node strength

Having characterized effects at the level of individual edges, we next assessed the impact of motion on node strength, a node-level measure that is used in many connectomic analyses to define network hubs. For each region in the 220-node parcellation, we estimated the cross-subject correlation between its strength (i.e., the sum of a nodes edge weights) and head motion of that person (henceforth termed QC-strength correlation). Figure 5 shows the proportion of nodes with a significant QC-strength correlation (*p* < .05, uncorrected) across different consistency thresholds. Pipelines using SSW edge weighting showed consistent effects of motion on node strength across different consistency thresholds: specifically, more stringent thresholds increased the proportion of nodes with a significant QC-strength correlation. The converse was observed specifically for pipelines using EDDY2, iFOD2, and FA weighting, where *less* stringent consistency thresholds *increased* the proportion of significant QC-Strength correlations.

**Figure 5.**
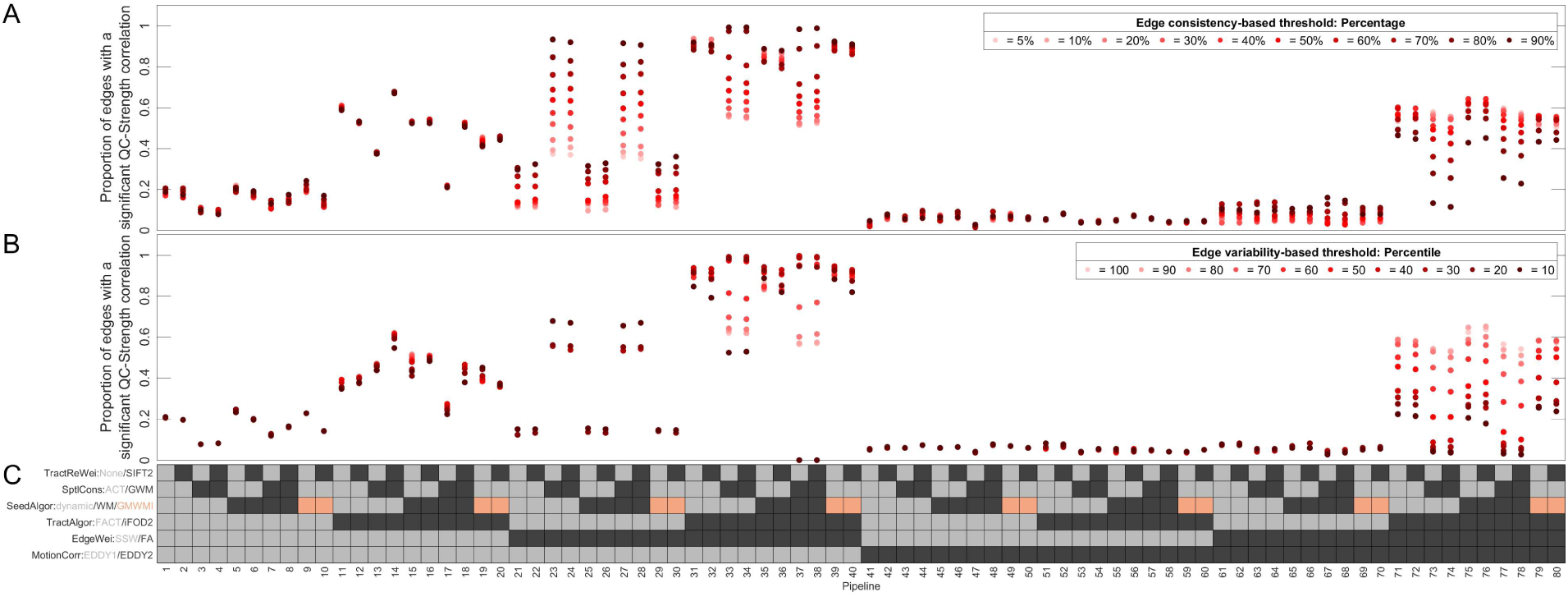
The effect of in-scanner motion (*ABS*_*all*_) on node strength when using a 220 node parcellation in 80 combinations of preprocessing choices across different edge consistency and variability thresholds. (**A**) The percentage of nodes (y-axis) which had a significant QC-strength correlation (y-axis) for each pipeline (x-axis) across edge consistency-based thresholds from 5% to 90%. (**B**) The percentage of nodes (y-axis) which had a significant QC-strength correlation (y-axis) for each pipeline (x-axis) across edge variability-based thresholds from the10^th^to 100^th^ percentile. In (**A**) and (**B**) darker colours indicate a more stringent threshold. (**C**) The preprocessing options used in each pipeline. Each row corresponds to a preprocessing step, with the possible options for that step colour-coded; the colour of the squares in each column indicate the specific methods used for a given pipeline.

As done previously (Baum et al., 2018), we used a 50% edge consistency-based threshold to calculate node strength and then assessed how node strength correlated with motion. The mean magnitude of correlations across pipelines varied from 0.13—0.31, with the strongest correlation of any node in any pipeline using the 220 node parcellation being -0.46. QC-strength correlations differed markedly across pipelines: in some pipelines, both positive and negative correlations could be observed (Figure 6A); whereas in others, widespread positive correlations were found (typically when iFOD2, SSW edge weighting, and EDDY1 were used; Figure 6B). With EDDY2, few nodes displayed any significant QC-strength relationships regardless of other aspects of the preprocessing pipeline (Figure 6D-E); the exception being when FA weighting and iFOD2 were used, where anatomically distributed and significant negative QC-strength correlations were observed with either EDDY1 (Figure 6C) or EDDY2 (Figure 6F), albeit with reduced magnitude in the latter case. Across pipelines, there was little consistency as to which regions showed significant QC-strength correlations, with no node exhibiting a significant correlation in more than 24% of pipelines (Fig S17).

**Figure 6.**
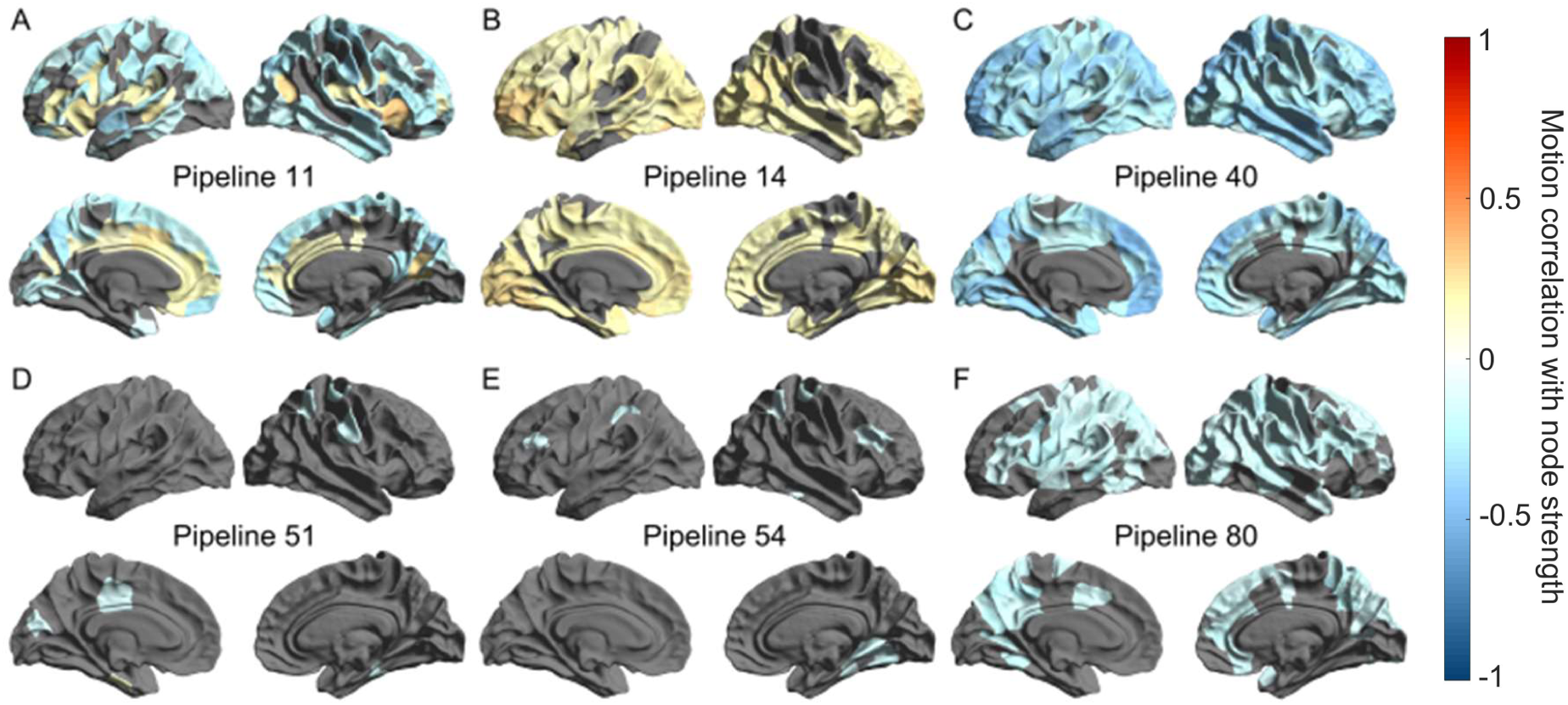
Correlation between in-scanner motion (quantified by mean absolute volume-to-volume displacement) and node strength for six different pipelines. Individual networks were thresholded using an edge-based consistency threshold of 50%, and each nodes’ strength across participants was correlated with mean absolute volume-to-volume displacement. Grey indicates a non-significant correlation. Pipeline numbers correspond to those in Figures 2, 3,5, and 7. (**A**) Significant QC-strength correlations for (**A**) iFOD2, SSW edge weighting, ACT mask, and with EDDY1(pipeline 11 shown as an example); (**B**) iFOD2, SSW edge weighting, GWM mask, and EDDY1 (pipeline 14 shown as an example); (**C**) iFOD2 and FA edge weighting, with EDDY1 (pipeline 40 shown as an example); (**D**) iFOD2, SSW edge weighting, ACT mask, and with EDDY2 (pipeline 51 shown as an example); (**E**) iFOD2, SSW edge weighting, GWM mask, with EDDY2 (pipeline 54 shown as an example); (**F**) iFOD2 and FA edge weighting, with EDDY2 (pipeline 80 shown as an example). The spatial topography, magnitude and polarity of QC-Strength correlations varies considerably across pipelines.

#### Node strength sequence variation across pipelines

Given the finding that motion can affect node strength estimates in a pipeline-dependent way, we sought to determine whether preprocessing pipelines influenced which nodes were defined as hubs; i.e. we sought to determine whether the relative ranking of nodes based on node strength was subject to variation, or whether it was preserved across pipelines. To do this, we calculated node strength rank (NSR) correlations (Spearman correlation of node strengths in a given pipeline) between every pair of pipelines. Finally, we reordered the NSR correlation matrix using hierarchical clustering (Figure 7).

**Figure 7.**
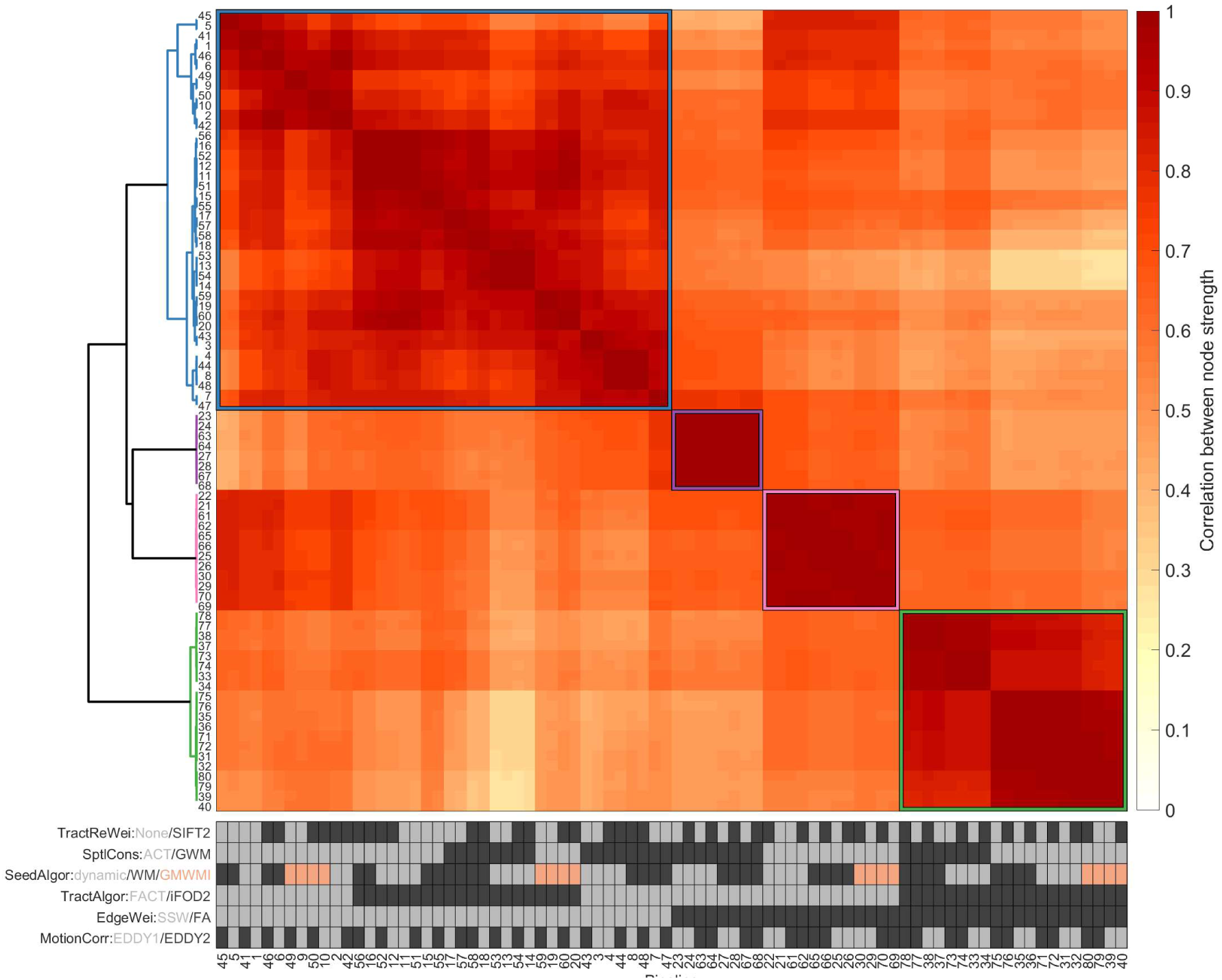
Node strength rank correlations. The correlation matrix has been reordered using hierarchical clustering (using correlation distances between rows and average linkage). Dendrogram on left, pipeline details at bottom. Four main clusters are apparent: pipelines using SSW edge weighting; pipelines using FA edge weighting, a GWM mask and FACT; pipelines using FA edge weighting, ACT and FACT; and pipelines using FA edge weighting and iFOD2.

This analysis revealed two key findings. First, there was considerable variability in NSR correlations between pairs of pipelines: 0.27<*ρ*<1.00. Thus, the spatial topography of a given node strength map, and therefore the designation of nodes as hubs, varied considerably depending on how the data were processed. Secondly, similarity in NSR was largely dictated by specific preprocessing steps. Figure 7 indicates the presence of four discrete clusters in the pipeline-by-pipeline correlation matrix. These clusters were predominantly determined by the tractography algorithm, streamline spatial constraints, and edge weighting methods used (note that these preprocessing steps almost entirely drove the clustering solution regardless of the edge consistency threshold being used). Pipelines that used SSW edge weighting grouped together in one cluster (blue cluster). NSR correlations between these pipelines ranging between 0.53 and 1.00 and were thus reasonably consistent. Other pipelines – all of which used FA edge weighting – were further stratified by the tractography algorithm and type of streamline spatial constraint being used. Specifically, pipelines using a GWM mask and FACT grouped together (purple cluster in dendrogram; 0.99<*ρ*<1.00), as did those using ACT and FACT (pink cluster in dendrogram; 0.64<*ρ*<1.00). Finally, pipelines using iFOD2 and FA were clustered together (green cluster in dendrogram; 0.80<*ρ*<1.00). We illustrate the two most dissimilar pipelines (*ρ* = 0.27) in Figure 8, where we mapped node strength across the brain for pipelines 39 and 53: with pipeline 39 (EDDY1, FACT, ACT, GWMI seeding, no SIFT and FA edge weighting), the hubs are located in parietal and posterior cingulate regions; with pipeline 53 (EDDY2, FACT, a GWM mask, dynamic seeding, no SIFT and SSW edge weighting), the hubs are located primarily in the occipital lobe. The most dissimilar pipelines (mean correlation of 0.277 ± 0.004) were those that used iFOD2, ACT, FA edge weighing, and GMWMI seeding (pipelines 13, 14, 53, 54), as compared to those which using iFOD2, a GWM mask, SSW edge weighting and dynamic seeding (pipelines 39, 40, 79, 80).

**Figure 8.**
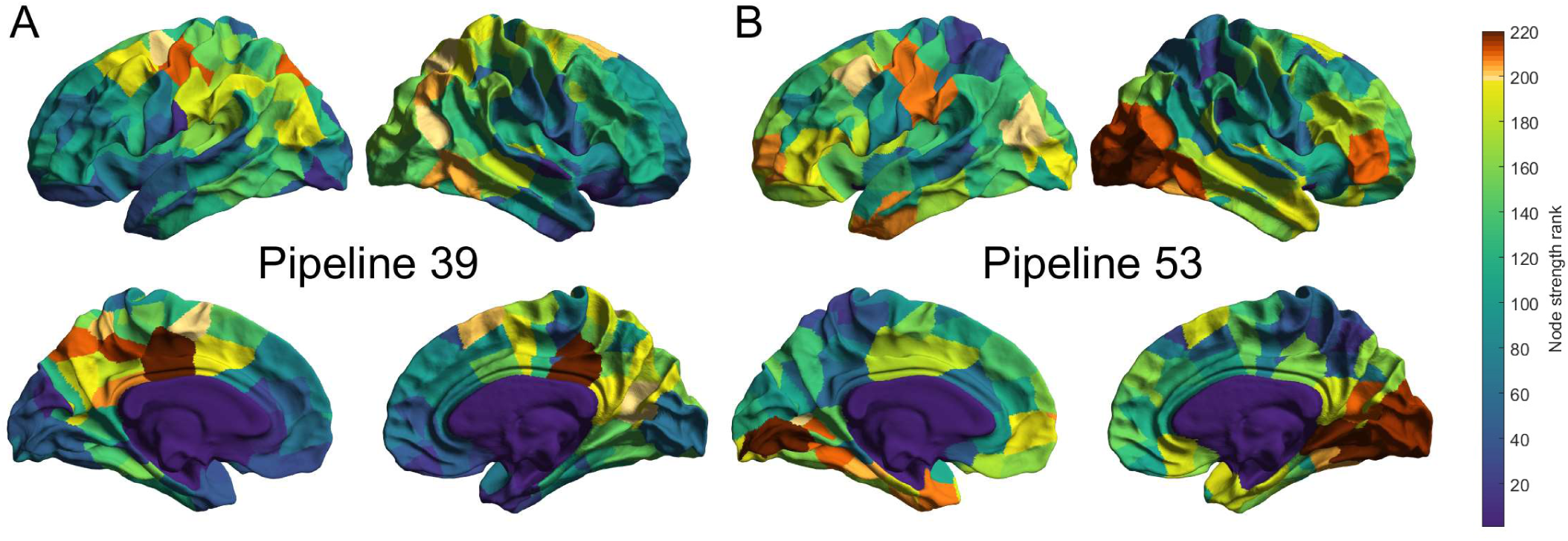
Spatial maps of mean node strength rankings for two dissimilar pipelines. A higher rank indicates the node has a higher strength. (**A**) Strength spatial distribution for pipeline 39, which used EDDY1, FACT, ACT, GWMI seeding, no SIFT and FA edge weighting. (**B**) Strength spatial distribution for pipeline 52, which used EDDY2, FACT, a GWM mask, dynamic seeding, no SIFT and SSW edge weighting.

## Discussion

Our experiments sought to characterise the degree to which different preprocessing choices mitigate motion-related contamination of diffusion MRI by examining 240 pipelines combining 16 different choices that are commonly made when preprocessing DWI data in preparation for connectome construction. Our analysis indicates that the degree of motion contamination varies considerably across pipelines. The biggest impact involved the use of state-of-the-art DWI preprocessing capabilities - specifically outlier replacement and within-volume motion-correction - which dramatically reduced the proportion of edges showing significant QC-SC correlations. Beyond this primary effect, the use of FA edge-weighting and iFOD2 resulted in higher QC-SC correlations. The strongest motion effects tended to be concentrated on long-range, inconsistent edges. Critically, these effects are distributed throughout the connectome in a pipeline-dependent manner. Motion effects can have a major impact on subsequent graph-theoretical analyses, leading to marked differences in conclusions, such as the labelling of which brain regions are network hubs.

### Outlier replacement and within-volume motion are powerful methods for mitigating motion-related artefact in diffusion MRI connectomics

Compared to EDDY1, the use of EDDY2 dramatically reduced the proportion of edges with significant QC-SC correlations. Our post-hoc analysis indicated that, of the novel capabilities incorporated in EDDY2, both outlier replacement and within-volume correction contribute to the superior efficacy of EDDY2 in this context, though the latter was particularly influential for FA-weighted connectomes. Incorporating outlier replacement and within-volume correction into connectome construction pipelines is therefore strongly encouraged as it confers significant advantages in mitigating motion-related contamination.

### The effect of tractography algorithm

The magnitude of QC-SC correlations was substantially affected by the choice of tractography algorithm. When using EDDY1, FACT resulted in fewer significant QC-SC correlations than iFOD2 (although this difference was greatly diminished when using EDDY2). This might be partly because algorithms like FACT have difficulty reconstructing complex fibre pathways, and these pathways may be more susceptible to motion related effects, rather than reflecting any intrinsic robustness to motion. Regardless, it is still useful to know which tractography algorithms may be more affected by motion as this can be one factor to consider when trying to decide on an algorithm to use, or when considering the application of other preprocessing steps.

### The effect of edge weighting

Connectomes in which edge weights were defined using the mean FA of the connecting bundle exhibited stronger QC-SC correlations than those constructed using SSW edge weighting, particularly when used in conjunction with EDDY1. Other studies have noted that FA is highly susceptible to contamination by motion (Jones & Basser, 2004; Le Bihan et al., 2006; Tijssen et al., 2009), and that FA-weighting increases the proportion of edges significantly correlated with motion estimates compared to SSW-weighting (Baum et al., 2018). This effect was diminished by the use of EDDY2, consistent with prior work showing that outlier-replacement and within-volume correction improves the measurement of FA in the presence of motion artefacts (Andersson et al., 2017, 2016).

For both FA and SSW edge weighting, inconsistent long-range connections displayed the strongest motion effects, although these effects were neither systematically positive nor negative. Long-range connections tend to pass through deep WM in addition to superficial WM areas, with the former having higher FA values/simpler FOD shapes, but they also inherit a greater number of opportunities to stray from the ideal reconstruction trajectory; this increases the prospect of motion-related image contamination biasing connectivity estimates in one direction or the other depending on the surrounding WM fibre geometry and the resulting erroneous trajectories.

However, because inconsistent edges by definition have fewer non-zero values to use in calculating QC-SC values, this absence of zero values may result in overinflated correlations with motion estimates. Indeed, when we included all edge weight values (i.e., did not exclude subjects whose edge weight was zero) in the estimation of QC-SC correlations, the magnitudes of these correlations decreased. Although the inclusion of zero edge weights did not have a major effect on the *proportion* of significant QC-SC correlations, it did change the relationships between QC-SC, edge consistency, length, and weight variability, becoming more uniform across the range of edge weights in line with previous results (Baum et al., 2018). Therefore, the way in which QC-SC is calculated (i.e., including or excluding zero edge weights) can alter the conclusions that one draws when assessing efficacy of motion contamination removal.

Although we favour excluding subjects for whom there is no edge when calculating this value (as otherwise the correlation becomes more reflective of edge *presence/absence* than actual variability in edge weight values), this omission of subject data must also be considered in its own right: a pipeline where connectome edges disappear entirely due to motion should not be considered “robust to motion” only because values for those subjects where the edge is preserved do not correlate with motion estimates.

### Interactions between preprocessing steps

Some preprocessing steps showed complex interactions between one another. For example, when using EDDY1, FA-weighted connectomes generated with ACT showed fewer significant QC-SC correlations than those generated with a GWM mask, whereas this trend was reversed for SSW-weighted connectomes (i.e. using ACT resulted in more significant QC-SC correlations than GWM masking). When using EDDY2, the combination of ACT, iFOD2, and FA weighting yielded higher residual motion contamination than when using a GWM mask, reversing the trend observed following EDDY1, albeit with a reduced magnitude of that difference (the mean difference in the proportion of significant QC-SC correlations between pipelines using ACT or not being 24.4% for EDDY1, 6.46% for EDDY2). These findings indicate that the combination of different preprocessing steps can interact to influence residual noise structure in the connectome data.

### Preprocessing choices influence network architecture

Our region-level analysis of the correlation between node strength (specifically rank thereof) and head motion revealed three noteworthy findings. First, as with the edge-level analysis, EDDY2 dramatically reduced QC-strength correlations. Second, the spatial distribution of these correlations varied considerably across pipelines, indicating that there were no regional ‘hotspots’ consistently affected by motion: the specific brain regions affected by motion depend strongly on the preprocessing choices made. Finally, node strength rankings, which generally determine which nodes are considered to be hubs, varied considerably across different pipelines. Variability was smaller between those pipelines using SSW (mean correlation = 0.83, *SD* = 0.11). Correlations between pipelines using FA-weighting were more heterogeneous (mean correlation = 0.71, *SD* = 0.19), and such pipelines were classified into three clusters where node strength rankings were internally consistent: one that combined iFOD2 with FA, and two that segregated FACT-based FA-weighted connectomes into those that used ACT or a grey-white matter mask. Preprocessing choices may thus have a significant impact on the final architecture of the connectome; as shown in Figure 8, nodes of high strength in visual cortex and rostral prefrontal cortex in one pipeline (panel B) have low strength in another pipeline (panel A). These variations result in very different hub locations in the connectome, and hence experimental conclusions that may be drawn from such. Although, the distinction between SSW-weighting and FA-weighting is known to result in connectomes with a very different distribution of weights, researchers must be cognisant of the strong dependence of experimental outcomes on the details of their reconstruction pipeline.

## Limitations

Our analyses focused on preprocessing steps implemented in two software packages: FSL and *MRtrix3*. Although these are among the most popular in connectomics, many other programs are available, each of which rely on different models, algorithms, and assumptions. We opted here to focus on well-defined choices available within integrated packages, rather than exploring the full parameter space of possible combinations between steps implemented in different packages. Our findings represent an initial step that can be extended by further exploration of other tools and algorithms.

We restricted our analyses to examining the fundamental decisions with respect to pipeline design, rather than the myriad tuneable parameters within each. For example, in tractography algorithms an FA or FOD amplitude threshold is often set so streamlines will terminate when they reach a voxel below that threshold. Given the vulnerability of FA to motion (Jones & Basser, 2004; Le Bihan et al., 2006; Tijssen et al., 2009), differences in this threshold may affect the extent to which residual motion artefacts bias tractography. From the few studies that have evaluated in detail how the tuning of tractography and diffusion parameters affects structural connectivity estimates, initial results suggest they can be consequential (Bastiani, Shah, Goebel, & Roebroeck, 2012; Li et al., 2012; Yeh et al., 2019). Given the large number of available diffusion and tractography approaches, it is important to understand the effect of these more fine-grained parameter choices, but these were beyond the feasible scope of our study.

Finally, one difficulty in evaluating the impact of motion on structural connectivity is the absence of a ground truth. This limitation makes it difficult to clearly establish the extent to which head motion is driving false-positive or false-negative connections. Previous studies examining the validity of different tractography algorithms have used synthetic DWI data where ground truth fibre bundles are known (Maier-Hein et al., 2017; Schilling, Daducci, et al., 2019; Schilling, Nath, et al., 2019), or have relied on comparisons to axonal tract-tracing data (Sinke et al., 2018; Thomas et al., 2014). Simulations of head motion in a model where ground truths are known could allow for a more detailed characterisation of how streamline reconstructions are affected under such conditions, which in turn would allow for a better understanding of how preprocessing steps are or are not sensitive to residual motion-related artefacts.

## Conclusions

Our findings indicate that the choice of preprocessing pipeline strongly influences the extent to which motion may contaminate estimates of structural connectivity. These effects are heterogeneous throughout the network, affecting different network elements in a pipeline-dependent way, and with complex interactions between pipeline components. The fact that the final topology and topography of the network is highly variable and depends on prior preprocessing choices is an important consideration when interpreting the results of connectomic analysis.

Importantly, the use of outlier replacement and within-volume motion correction (as implemented in EDDY2) can dramatically mitigate the residual effects of motion on connectome construction; we therefore encourage the use of this approach in future dMRI studies.

## Code and data availability

All the data used in this study is openly available online on Figshare at https://figshare.com/s/3310385f29a156c93ca3. Scripts to analyse this data are available on gitHub at https://github.com/BMHLab/MotionStructuralConnectivity.

## Reference list

Anderson, A. W., & Gore, J. C. (1994). Analysis and correction of motion artifacts in diffusion weighted imaging. Magnetic Resonance in Medicine, 32(3), 379–387. https://doi.org/10.1002/mrm.1910320313

Andersson, J. L. R., Graham, M. S., Drobnjak, I., Zhang, H., Filippini, N., & Bastiani, M. (2017). Towards a comprehensive framework for movement and distortion correction of diffusion MR images: Within volume movement. NeuroImage, 152(February), 450–466. https://doi.org/10.1016/j.neuroimage.2017.02.085

Andersson, J. L. R., Graham, M. S., Zsoldos, E., & Sotiropoulos, S. N. (2016). Incorporating outlier detection and replacement into a non-parametric framework for movement and distortion correction of diffusion MR images. NeuroImage, 141, 556–572. https://doi.org/10.1016/j.neuroimage.2016.06.058

Andersson, J. L. R., & Skare, S. (2002). A model-based method for retrospective correction of geometric distortions in diffusion-weighted EPI. NeuroImage, 16(1), 177–199. https://doi.org/10.1006/nimg.2001.1039

Andersson, J. L. R., Skare, S., & Ashburner, J. (2003). How to correct susceptibility distortions in spinecho echo-planar images: Application to diffusion tensor imaging. NeuroImage, 20(2), 870–888. https://doi.org/10.1016/S1053-8119(03)00336-7

Andersson, J. L. R., & Sotiropoulos, S. N. (2016). An integrated approach to correction for off-resonance effects and subject movement in diffusion MR imaging. NeuroImage, 125, 1063–1078. https://doi.org/10.1016/j.neuroimage.2015.10.019

Baker, S. T. E., Lubman, D. I., Yucel, M., Allen, N. B., Whittle, S., Fulcher, B. D., … Fornito, A. (2015). Developmental Changes in Brain Network Hub Connectivity in Late Adolescence. Journal of Neuroscience, 35(24), 9078–9087. https://doi.org/10.1523/JNEUROSCI.5043-14.2015

Bassett, D. S., Xia, C. H., & Satterthwaite, T. D. (2018). Understanding the Emergence of Neuropsychiatric Disorders with Network Neuroscience. Biological Psychiatry: Cognitive Neuroscience and Neuroimaging. https://doi.org/10.1016/j.bpsc.2018.03.015

Bastiani, M., Cottaar, M., Fitzgibbon, S. P., Suri, S., Alfaro-Almagro, F., Sotiropoulos, S. N., … Andersson, J. L. R. (2019). Automated quality control for within and between studies diffusion MRI data using a non-parametric framework for movement and distortion correction. NeuroImage, 184(May 2018), 801–812. https://doi.org/10.1016/j.neuroimage.2018.09.073

Bastiani, M., Shah, N. J., Goebel, R., & Roebroeck, A. (2012). Human cortical connectome reconstruction from diffusion weighted MRI: The effect of tractography algorithm. NeuroImage, 62(3), 1732–1749. https://doi.org/10.1016/j.neuroimage.2012.06.002

Baum, G. L., Ciric, R., Roalf, D. R., Betzel, R. F., Moore, T. M., Shinohara, R. T., … Satterthwaite, T. D. (2017). Modular Segregation of Structural Brain Networks Supports the Development of Executive Function in Youth. Current Biology, 27(11), 1561-1572.e8. https://doi.org/10.1016/j.cub.2017.04.051

Baum, G. L., Roalf, D. R., Cook, P. A., Ciric, R., Rosen, A. F. G., Xia, C., … Satterthwaite, T. D. (2018). The impact of in-scanner head motion on structural connectivity derived from diffusion MRI. NeuroImage, 173(November 2017), 275–286. https://doi.org/10.1016/j.neuroimage.2018.02.041

Beaulieu, C. (2002). The basis of anisotropic water diffusion in the nervous system - A technical review. NMR in Biomedicine, 15(7–8), 435–455. https://doi.org/10.1002/nbm.782

Behrens, T. E. J., Johansen-Berg, H., Woolrich, M. W., Smith, S. M., Wheeler-Kingshott, C. A. M., Boulby, P. A., … Matthews, P. M. (2003). Non-invasive mapping of connections between human thalamus and cortex using diffusion imaging. Nature Neuroscience, 6(7), 750–757. https://doi.org/10.1038/nn1075

Behrens, T. E. J., Woolrich, M. W., Jenkinson, M., Johansen-Berg, H., Nunes, R. G., Clare, S., … Smith, S. M. (2003). Characterization and Propagation of Uncertainty in Diffusion-Weighted MR Imaging. Magnetic Resonance in Medicine, 50(5), 1077–1088. https://doi.org/10.1002/mrm.10609

Bullmore, E. T., & Sporns, O. (2009). Complex brain networks: graph theoretical analysis of structural and functional systems. Nature Reviews Neuroscience, 10(3), 186–198. https://doi.org/10.1038/nrn2575

Bullmore, E. T., & Sporns, O. (2012). The economy of brain network organization. Nature Reviews Neuroscience, 13(5), 336–349. https://doi.org/10.1038/nrn3214

Cao, M., Huang, H., & He, Y. (2017). Developmental Connectomics from Infancy through Early Childhood. Trends in Neurosciences, 40(8), 494–506. https://doi.org/10.1016/j.tins.2017.06.003

Ciric, R., Wolf, D. H., Power, J. D., Roalf, D. R., Baum, G. L., Ruparel, K., … Satterthwaite, T. D. (2017). Benchmarking of participant-level confound regression strategies for the control of motion artifact in studies of functional connectivity. NeuroImage, 154(March), 174–187. https://doi.org/10.1016/j.neuroimage.2017.03.020

Crossley, N. A., Mechelli, A., Scott, J., Carletti, F., Fox, P. T., Mcguire, P., & Bullmore, E. T. (2014). The hubs of the human connectome are generally implicated in the anatomy of brain disorders. Brain, 137(8), 2382–2395. https://doi.org/10.1093/brain/awu132

Daducci, A., Dal Palù, A., Lemkaddem, A., & Thiran, J. P. (2015). COMMIT: Convex optimization modeling for microstructure informed tractography. IEEE Transactions on Medical Imaging, 34(1), 246–257. https://doi.org/10.1109/TMI.2014.2352414

de Reus, M. a., & van den Heuvel, M. P. (2013). Estimating false positives and negatives in brain networks. NeuroImage, 70, 402–409. https://doi.org/10.1016/j.neuroimage.2012.12.066

Desikan, R. S., Ségonne, F., Fischl, B., Quinn, B. T., Dickerson, B. C., Blacker, D., … Killiany, R. J. (2006). An automated labeling system for subdividing the human cerebral cortex on MRI scans into gyral based regions of interest. NeuroImage, 31(3), 968–980. https://doi.org/10.1016/j.neuroimage.2006.01.021

Fair, D. A., Nigg, J. T., Iyer, S., Bathula, D., Mills, K. L., Dosenbach, N. U. F., … Milham, M. P. (2013). Distinct neural signatures detected for ADHD subtypes after controlling for micro-movements in resting state functional connectivity MRI data. Frontiers in Systems Neuroscience, 6(February), 1–31. https://doi.org/10.3389/fnsys.2012.00080

Fischl, B. (2012). FreeSurfer. NeuroImage, 62(2), 774–781. https://doi.org/10.1016/j.neuroimage.2012.01.021

Fischl, B., Salat, D. H., Busa, E., Albert, M., Dieterich, M., Haselgrove, C., … Dale, A. M. (2002). Whole brain segmentation: Automated labeling of neuroanatomical structures in the human brain. Neuron, 33(3), 341–355. https://doi.org/10.1016/S0896-6273(02)00569-X

Fornito, A., Zalesky, A., Bassett, D. S., Meunier, D., Ellison-Wright, I., Yucel, M., … Bullmore, E. T. (2011). Genetic Influences on Cost-Efficient Organization of Human Cortical Functional Networks. Journal of Neuroscience, 31(9), 3261–3270. https://doi.org/10.1523/JNEUROSCI.4858-10.2011

Fornito, A., Zalesky, A., & Breakspear, M. (2015). The connectomics of brain disorders. Nature Reviews Neuroscience, 16(3), 159–172. https://doi.org/10.1038/nrn3901

Fornito, A., Zalesky, A., & Bullmore, E. T. (2010). Network scaling effects in graph analytic studies of human resting-state fMRI data. Frontiers in Systems Neuroscience, 4(June), 1–16. https://doi.org/10.3389/fnsys.2010.00022

Fornito, A., Zalesky, A., & Bullmore, E. T. (2016). Fundamentals of Brain Network Analysis. London: Academic Press.

Girard, G., Whittingstall, K., Deriche, R., & Descoteaux, M. (2014). Towards quantitative connectivity analysis: Reducing tractography biases. NeuroImage, 98, 266–278. https://doi.org/10.1016/j.neuroimage.2014.04.074

Glasser, M. F., Coalson, T. S., Robinson, E. C., Hacker, C. D., Harwell, J., Yacoub, E., … Van Essen, D. C. (2016). A multi-modal parcellation of human cerebral cortex. Nature, 536(7615), 171–178. https://doi.org/10.1038/nature18933

Goñi, J., van den Heuvel, M. P., Avena-Koenigsberger, A., Velez de Mendizabal, N., Betzel, R. F., Griffa, A., … Sporns, O. (2014). Resting-brain functional connectivity predicted by analytic measures of network communication. Proceedings of the National Academy of Sciences of the United States of America, 111(2), 833–838. https://doi.org/10.1073/pnas.1315529111

Goscinski, W. J., McIntosh, P., Felzmann, U., Maksimenko, A., Hall, C. J., Gureyev, T., … Egan, G. F. (2014). The multi-modal Australian ScienceS imaging and visualization environment (MASSIVE) high performance computing infrastructure: Applications in neuroscience and neuroinformatics research. Frontiers in Neuroinformatics, 8(MAR), 1–13. https://doi.org/10.3389/fninf.2014.00030

Greve, D. N., & Fischl, B. (2009). Accurate and robust brain image alignment using boundary-based registration. NeuroImage, 48(1), 63–72. https://doi.org/10.1016/j.neuroimage.2009.06.060

Honey, C. J., Thivierge, J. P., & Sporns, O. (2010). Can structure predict function in the human brain? NeuroImage, 52(3), 766–776. https://doi.org/10.1016/j.neuroimage.2010.01.071

Jenkinson, M., Bannister, P., Brady, M., & Smith, S. (2002). Improved Optimization for the Robust and Accurate Linear Registration and Motion Correction of Brain Images. NeuroImage, 17(2), 825–841. https://doi.org/10.1006/nimg.2002.1132

Jenkinson, M., Beckmann, C. F., Behrens, T. E. J., Woolrich, M. W., & Smith, S. M. (2012). FSL. NeuroImage, 62(2), 782–790. https://doi.org/10.1016/j.neuroimage.2011.09.015

Jenkinson, M., & Smith, S. (2001). A global optimisation method for robust affine registration of brain images. Medical Image Analysis, 5(2), 143–156. https://doi.org/10.1016/S1361-8415(01)00036-6

Jeurissen, B., Descoteaux, M., Mori, S., & Leemans, A. (2019). Diffusion MRI fiber tractography of the brain. NMR in Biomedicine, 32(4), e3785. https://doi.org/10.1002/nbm.3785

Jones, D. K. (2010). Challenges and limitations of quantifying brain connectivity in vivo with diffusion MRI. Imaging in Medicine, 2(3), 341–355. https://doi.org/10.2217/iim.10.21

Jones, D. K., & Basser, P. J. (2004). “Squashing peanuts and smashing pumpkins”: How noise distorts diffusion-weighted MR data. Magnetic Resonance in Medicine, 52(5), 979–993. https://doi.org/10.1002/mrm.20283

Jones, D. K., Knösche, T. R., & Turner, R. (2013). White matter integrity, fiber count, and other fallacies: The do’s and don’ts of diffusion MRI. NeuroImage, 73, 239–254. https://doi.org/10.1016/j.neuroimage.2012.06.081

Le Bihan, D., Poupon, C., Amadon, A., & Lethimonnier, F. (2006). Artifacts and pitfalls in diffusion MRI. Journal of Magnetic Resonance Imaging, 24(3), 478–488. https://doi.org/10.1002/jmri.20683

Li, L., Rilling, J. K., Preuss, T. M., Glasser, M. F., & Hu, X. (2012). The effects of connection reconstruction method on the interregional connectivity of brain networks via diffusion tractography. Human Brain Mapping, 33(8), 1894–1913. https://doi.org/10.1002/hbm.21332

Ling, J., Merideth, F., Caprihan, A., Pena, A., Teshiba, T., & Mayer, A. R. (2012). Head injury or head motion? Assessment and quantification of motion artifacts in diffusion tensor imaging studies. Human Brain Mapping, 33(1), 50–62. https://doi.org/10.1002/hbm.21192

Liu, B., Zhu, T., & Zhong, J. (2015). Comparison of quality control software tools for diffusion tensor imaging. Magnetic Resonance Imaging, 33(3), 276–285. https://doi.org/10.1016/j.mri.2014.10.011

Maier-Hein, K. H., Neher, P. F., Houde, J.-C., Côté, M.-A., Garyfallidis, E., Zhong, J., … Descoteaux, M. (2017). The challenge of mapping the human connectome based on diffusion tractography. Nature Communications, 8(1), 1349. https://doi.org/10.1038/s41467-017-01285-x

Morgan, S. E., White, S. R., Bullmore, E. T., & Vértes, P. E. (2018). A network neuroscience approach to typical and atypical brain development. Biological Psychiatry: Cognitive Neuroscience and Neuroimaging. https://doi.org/10.1016/j.bpsc.2018.03.003

Mori, S., Crain, B. J., Chacko, V. P., & van Zijl, P. C. (1999). Three-dimensional tracking of axonal projections in the brain by magnetic resonance imaging. Annals of Neurology, 45(2), 265–269. https://doi.org/10.1002/1531-8249(199902)45:2<265::AID-ANA21>3.0.CO;2-3

Mori, S., & van Zijl, P. C. M. (2002). Fiber tracking: principles and strategies - a technical review. NMR in Biomedicine, 15(7–8), 468–480. https://doi.org/10.1002/nbm.781

Oguz, I., Gerig, G., Johnson, H. J., Farzinfar, M., Styner, M., Matsui, J., … Budin, F. (2014). DTIPrep: quality control of diffusion-weighted images. Frontiers in Neuroinformatics, 8(January), 1–11. https://doi.org/10.3389/fninf.2014.00004

Oldham, S., & Fornito, A. (2019). The development of brain network hubs. Developmental Cognitive Neuroscience, 36. https://doi.org/10.1016/j.dcn.2018.12.005

Oldham, S., Fulcher, B., Parkes, L., Arnatkevičiūtė, A., Suo, C., & Fornito, A. (2019). Consistency and differences between centrality measures across distinct classes of networks. Plos One, 14(7), e0220061. https://doi.org/10.1371/journal.pone.0220061

Parkes, L., Fulcher, B., Yücel, M., & Fornito, A. (2018). An evaluation of the efficacy, reliability, and sensitivity of motion correction strategies for resting-state functional MRI. NeuroImage, 171(December 2017), 415–436. https://doi.org/10.1016/j.neuroimage.2017.12.073

Patenaude, B., Smith, S. M., Kennedy, D. N., & Jenkinson, M. (2011). A Bayesian model of shape and appearance for subcortical brain segmentation. NeuroImage, 56(3), 907–922. https://doi.org/10.1016/j.neuroimage.2011.02.046

Pestilli, F., Yeatman, J. D., Rokem, A., Kay, K. N., & Wandell, B. A. (2014). Evaluation and statistical inference for human connectomes. Nature Methods, 11(10), 1058–1063. https://doi.org/10.1038/nmeth.3098

Power, J. D., Barnes, K. A., Snyder, A. Z., Schlaggar, B. L., & Petersen, S. E. (2012). Spurious but systematic correlations in functional connectivity MRI networks arise from subject motion. NeuroImage, 59(3), 2142–2154. https://doi.org/10.1016/j.neuroimage.2011.10.018

Reveley, C., Seth, A. K., Pierpaoli, C., Silva, A. C., Yu, D., Saunders, R. C., … Ye, F. Q. (2015). Superficial white matter fiber systems impede detection of long-range cortical connections in diffusion MR tractography. Proceedings of the National Academy of Sciences, 201418198. https://doi.org/10.1073/pnas.1418198112

Roalf, D. R., Quarmley, M., Elliott, M. A., Satterthwaite, T. D., Vandekar, S. N., Ruparel, K., … Gur, R. E. (2016). The impact of quality assurance assessment on diffusion tensor imaging outcomes in a large-scale population-based cohort. NeuroImage, 125, 903–919. https://doi.org/10.1016/j.neuroimage.2015.10.068

Roberts, J. A., Perry, A., Roberts, G., Mitchell, P. B., & Breakspear, M. (2017). Consistency-based thresholding of the human connectome. NeuroImage, 145, 118–129. https://doi.org/10.1016/j.neuroimage.2016.09.053

Rohde, G. K., Barnett, A. S., Basser, P. J., Marenco, S., & Pierpaoli, C. (2004). Comprehensive approach for correction of motion and distortion in diffusion-weighted MRI. Magnetic Resonance in Medicine, 51(1), 103–114. https://doi.org/10.1002/mrm.10677

Rubinov, M., & Sporns, O. (2010). Complex network measures of brain connectivity: Uses and interpretations. NeuroImage, 52(3), 1059–1069. https://doi.org/10.1016/j.neuroimage.2009.10.003

Sabaroedin, K., Tiego, J., Parkes, L., Sforazzini, F., Finlay, A., Johnson, B., … Fornito, A. (2019). Functional Connectivity of Corticostriatal Circuitry and Psychosis-like Experiences in the General Community. Biological Psychiatry, 86(1), 16–24. https://doi.org/10.1016/j.biopsych.2019.02.013

Satterthwaite, T. D., Elliott, M. A., Gerraty, R. T., Ruparel, K., Loughead, J., Calkins, M. E., … Wolf, D. H. (2013). An Improved Framework for Confound Regression and Filtering for Control of Motion Artifact in the Preprocessing of Resting-State Functional Connectivity Data. NeuroImage, 64, 240–256. https://doi.org/10.1016/j.neuroimage.2012.08.052

Satterthwaite, T. D., Wolf, D. H., Loughead, J., Ruparel, K., Elliott, M. A., Hakonarson, H., … Gur, R. E. (2012). Impact of in-scanner head motion on multiple measures of functional connectivity: Relevance for studies of neurodevelopment in youth. NeuroImage, 60(1), 623–632. https://doi.org/10.1016/j.neuroimage.2011.12.063

Schilling, K. G., Daducci, A., Maier-Hein, K., Poupon, C., Houde, J. C., Nath, V., … Descoteaux, M. (2019). Challenges in diffusion MRI tractography – Lessons learned from international benchmark competitions. Magnetic Resonance Imaging, 57(October 2018), 194–209. https://doi.org/10.1016/j.mri.2018.11.014

Schilling, K. G., Nath, V., Hansen, C., Parvathaneni, P., Blaber, J., Gao, Y., … Landman, B. A. (2019). Limits to anatomical accuracy of diffusion tractography using modern approaches. NeuroImage, 185(October 2018), 1–11. https://doi.org/10.1016/j.neuroimage.2018.10.029

Sinke, M. R. T., Otte, W. M., Christiaens, D., Schmitt, O., Leemans, A., van der Toorn, A., … Dijkhuizen, R. M. (2018). Diffusion MRI-based cortical connectome reconstruction: dependency on tractography procedures and neuroanatomical characteristics. Brain Structure and Function, 223(5), 2269–2285. https://doi.org/10.1007/s00429-018-1628-y

Skudlarski, P., Jagannathan, K., Calhoun, V. D., Hampson, M., Skudlarska, B. A., & Pearlson, G. (2008). Measuring brain connectivity: Diffusion tensor imaging validates resting state temporal correlations. NeuroImage, 43(3), 554–561. https://doi.org/10.1016/j.neuroimage.2008.07.063

Smith, R. E., Tournier, J. D., Calamante, F., & Connelly, A. (2012). Anatomically-constrained tractography: Improved diffusion MRI streamlines tractography through effective use of anatomical information. NeuroImage, 62(3), 1924–1938. https://doi.org/10.1016/j.neuroimage.2012.06.005

Smith, R. E., Tournier, J. D., Calamante, F., & Connelly, A. (2013). SIFT: Spherical-deconvolution informed filtering of tractograms. NeuroImage, 67, 298–312. https://doi.org/10.1016/j.neuroimage.2012.11.049

Smith, R. E., Tournier, J. D., Calamante, F., & Connelly, A. (2015a). SIFT2: Enabling dense quantitative assessment of brain white matter connectivity using streamlines tractography. NeuroImage, 119, 338–351. https://doi.org/10.1016/j.neuroimage.2015.06.092

Smith, R. E., Tournier, J. D., Calamante, F., & Connelly, A. (2015b). The effects of SIFT on the reproducibility and biological accuracy of the structural connectome. NeuroImage, 104, 253–265. https://doi.org/10.1016/j.neuroimage.2014.10.004

Smith, S. M. (2002). Fast robust automated brain extraction. Human Brain Mapping, 17(3), 143–155. https://doi.org/10.1002/hbm.10062

Smith, S. M., Jenkinson, M., Woolrich, M. W., Beckmann, C. F., Behrens, T. E. J., Johansen-Berg, H., … Matthews, P. M. (2004). Advances in functional and structural MR image analysis and implementation as FSL. NeuroImage, 23(SUPPL. 1), 208–219. https://doi.org/10.1016/j.neuroimage.2004.07.051

Sporns, O., Tononi, G., & Kötter, R. (2005). The human connectome: A structural description of the human brain. PLoS Computational Biology, 1(4), 0245–0251. https://doi.org/10.1371/journal.pcbi.0010042

Stam, C. J. (2014). Modern network science of neurological disorders. Nature Reviews Neuroscience, 15(10), 683–695. https://doi.org/10.1038/nrn3801

Thomas, C., Ye, F. Q., Irfanoglu, M. O., Modi, P., Saleem, K. S., Leopold, D. a., & Pierpaoli, C. (2014). Anatomical accuracy of brain connections derived from diffusion MRI tractography is inherently limited. Proceedings of the National Academy of Sciences, 111(46), 16574–16579. https://doi.org/10.1073/pnas.1405672111

Tijssen, R. H. N., Jansen, J. F. A., & Backes, W. H. (2009). Assessing and minimizing the effects of noise and motion in clinical DTI at 3 T. Human Brain Mapping, 30(8), 2641–2655. https://doi.org/10.1002/hbm.20695

Tournier, J. D., Calamante, F., & Connelly, A. (2007). Robust determination of the fibre orientation distribution in diffusion MRI: Non-negativity constrained super-resolved spherical deconvolution. NeuroImage, 35(4), 1459–1472. https://doi.org/10.1016/j.neuroimage.2007.02.016

Tournier, J. D., Calamante, F., & Connelly, A. (2010). Improved probabilistic streamlines tractography by 2nd order integration over fibre orientation distributions. Proceedings of the International Society for Magnetic Resonance in Medicine, 1670.

Tournier, J. D., Calamante, F., & Connelly, A. (2012). MRtrix: Diffusion tractography in crossing fiber regions. International Journal of Imaging Systems and Technology, 22(1), 53–66. https://doi.org/10.1002/ima.22005

Tournier, J. D., Smith, R., Raffelt, D., Tabbara, R., Dhollander, T., Pietsch, M., … Connelly, A. (2019). MRtrix3: A fast, flexible and open software framework for medical image processing and visualisation. NeuroImage, 116137. https://doi.org/10.1016/j.neuroimage.2019.116137

Tziortzi, A. C., Haber, S. N., Searle, G. E., Tsoumpas, C., Long, C. J., Shotbolt, P., … Gunn, R. N. (2014). Connectivity-based functional analysis of dopamine release in the striatum using diffusion-weighted MRI and positron emission tomography. Cerebral Cortex, 24(5), 1165–1177. https://doi.org/10.1093/cercor/bhs397

van den Heuvel, M. P., & Sporns, O. (2011). Rich-club organization of the human connectome. The Journal of Neuroscience : The Official Journal of the Society for Neuroscience, 31(44), 15775–15786. https://doi.org/10.1523/JNEUROSCI.3539-11.2011

van den Heuvel, M. P., & Sporns, O. (2013). Network hubs in the human brain. Trends in Cognitive Sciences, 17(12), 683–696. https://doi.org/10.1016/j.tics.2013.09.012

Veraart, J., Sijbers, J., Sunaert, S., Leemans, A., & Jeurissen, B. (2013). Weighted linear least squares estimation of diffusion MRI parameters: Strengths, limitations, and pitfalls. NeuroImage, 81, 335–346. https://doi.org/10.1016/j.neuroimage.2013.05.028

Yeh, C. H., Smith, R. E., Dhollander, T., Calamante, F., & Connelly, A. (2019). Connectomes from streamlines tractography: Assigning streamlines to brain parcellations is not trivial but highly consequential. NeuroImage, 199(April), 160–171. https://doi.org/10.1016/j.neuroimage.2019.05.005

Yendiki, A., Koldewyn, K., Kakunoori, S., Kanwisher, N., & Fischl, B. (2014). Spurious group differences due to head motion in a diffusion MRI study. NeuroImage, 88, 79–90. https://doi.org/10.1016/j.neuroimage.2013.11.027

Zalesky, A., Fornito, A., Cocchi, L., Gollo, L. L., van den Heuvel, M. P., & Breakspear, M. (2016). Connectome sensitivity or specificity: which is more important? NeuroImage, 142, 407–420. https://doi.org/10.1016/j.neuroimage.2016.06.035

Zalesky, A., Fornito, A., Harding, I. H., Cocchi, L., Yücel, M., Pantelis, C., & Bullmore, E. T. (2010). Whole-brain anatomical networks: Does the choice of nodes matter? NeuroImage, 50(3), 970–983. https://doi.org/10.1016/j.neuroimage.2009.12.027

Zhang, Y., Brady, M., & Smith, S. M. (2001). Segmentation of brain MR images through a hidden Markov random field model and the expectation-maximization algorithm. IEEE Transactions on Medical Imaging, 20(1), 45–57. https://doi.org/10.1109/42.906424

Zhao, T., Xu, Y., & He, Y. (2019). Graph theoretical modeling of baby brain networks. NeuroImage, 185(June), 711–727. https://doi.org/10.1016/j.neuroimage.2018.06.038

