## SupplementaryInfo for "The efficacy of different preprocessing steps in reducing motion-related confounds in diffusion MRI connectomics"

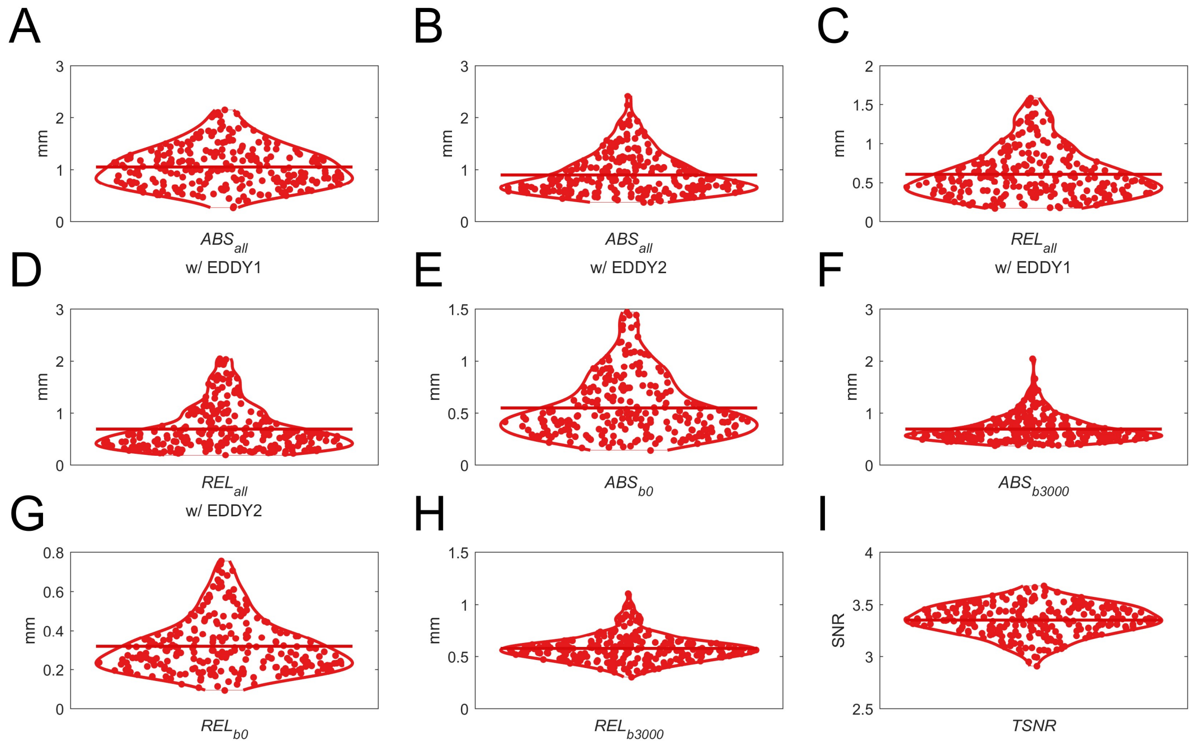


**Figure S1. Distribution of scores for different head motion measurements.**

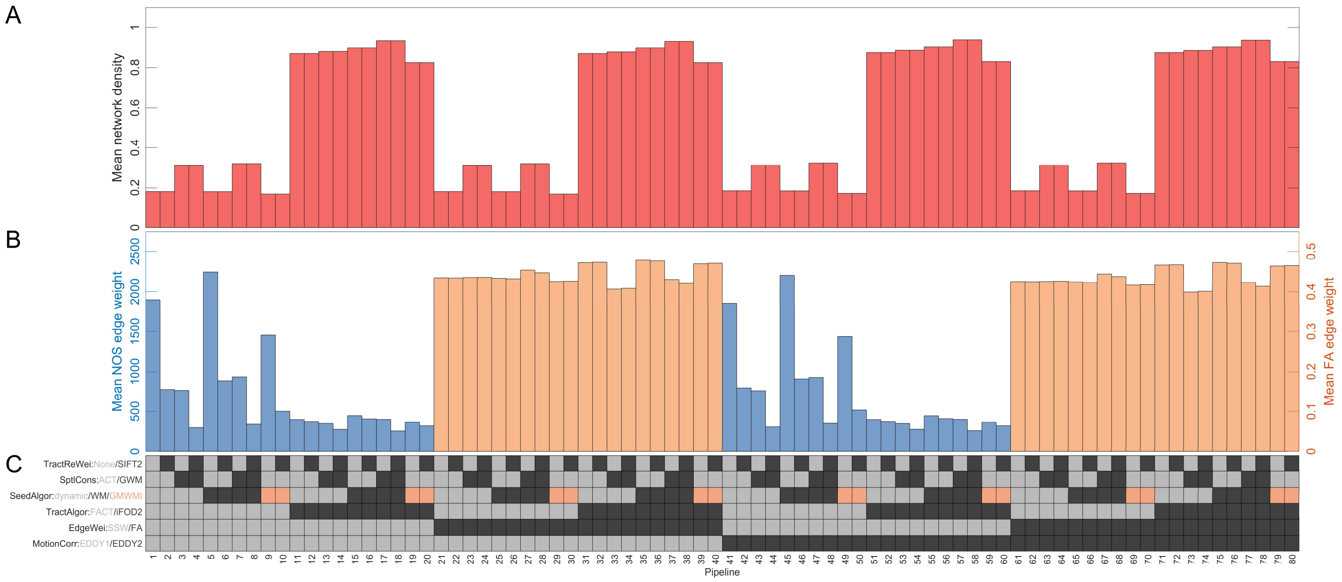


**Figure S2. Network properties in an 82 node parcellation in 80 combinations of processing choices.** (**A**) The mean participant network density (y-axis) in each pipeline (x-axis). (**B**) The mean edge weight for NOS (left y-axis) and FA (right y-axis) in each pipeline (x-axis). (**C**) The preprocessing options used in each pipeline. Each row corresponds to a preprocessing step, with the possible options for that step colour-coded; the colour of the squares in each column indicate the specific methods used for a given pipeline.

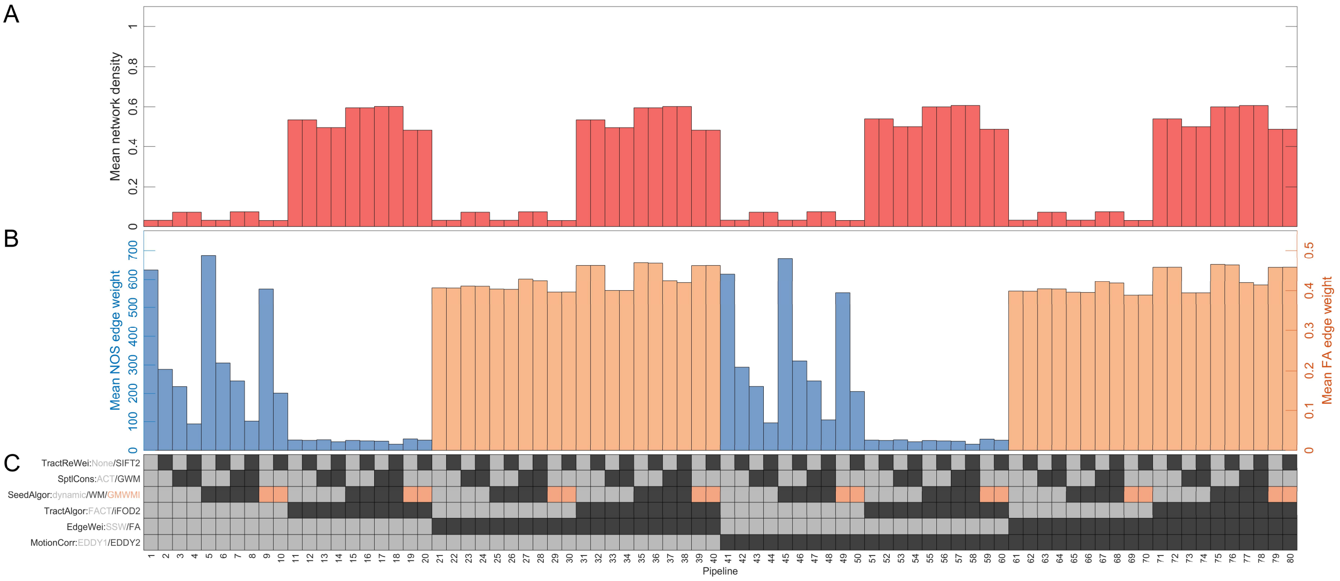


**Figure S3. Network properties in a 380 node parcellation in 80 combinations of processing choices.** (**A**) The mean participant network density (y-axis) in each pipeline (x-axis). (**B**) The mean edge weight for NOS (left y-axis) and FA (right y-axis) in each pipeline (x-axis). (**C**) The preprocessing options used in each pipeline. Each row corresponds to a preprocessing step, with the possible options for that step colour-coded; the colour of the squares in each column indicate the specific methods used for a given pipeline.

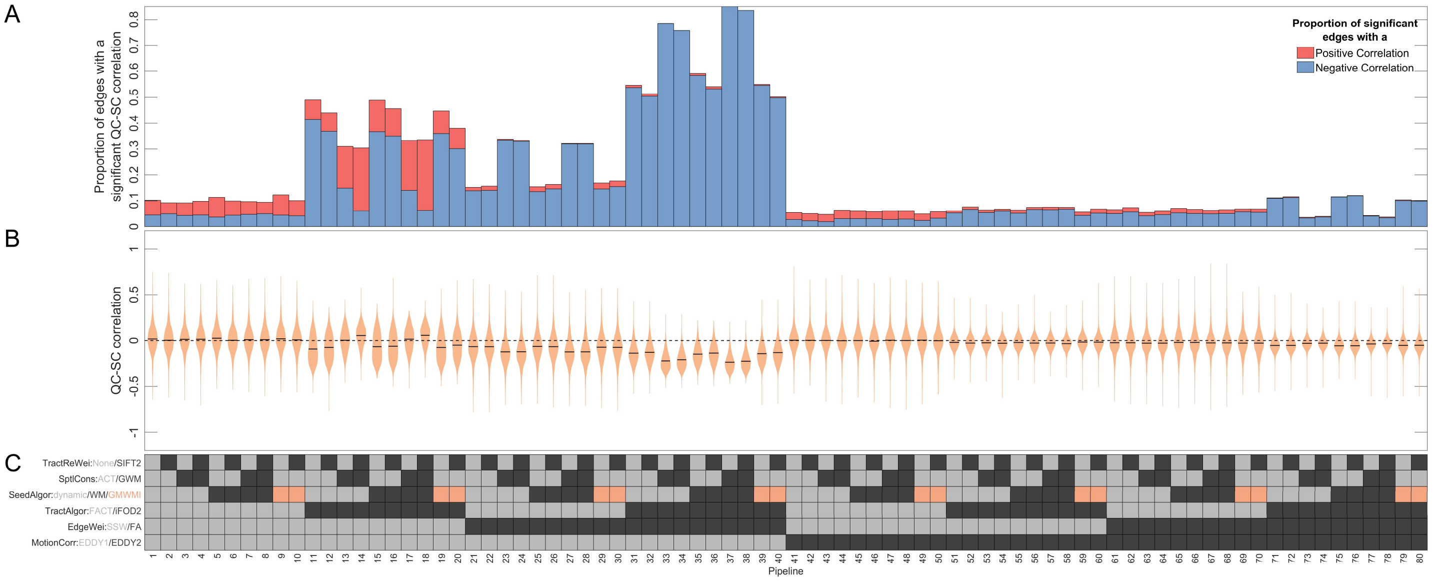


**Figure S4. The effect of in-scanner motion (*$ABS_{all}$*) on structural connectivity when using an 82 node parcellation.** (**A**) The proportion of edges (y-axis) that had a significant (*p* < .05, uncorrected) QC-SC correlation in each pipeline (x-axis). Each bar is coloured to show out of those significant edges, what proportion were negative (blue) or positive (red). (**B**) The full distributions of QC-SC correlations (y-axis) for each pipeline (x-axis). (**C**) The preprocessing options used in each pipeline. Each row corresponds to a preprocessing step, with the possible options for that step colour-coded; the colour of the squares in each column indicate the specific methods used for a given pipeline.

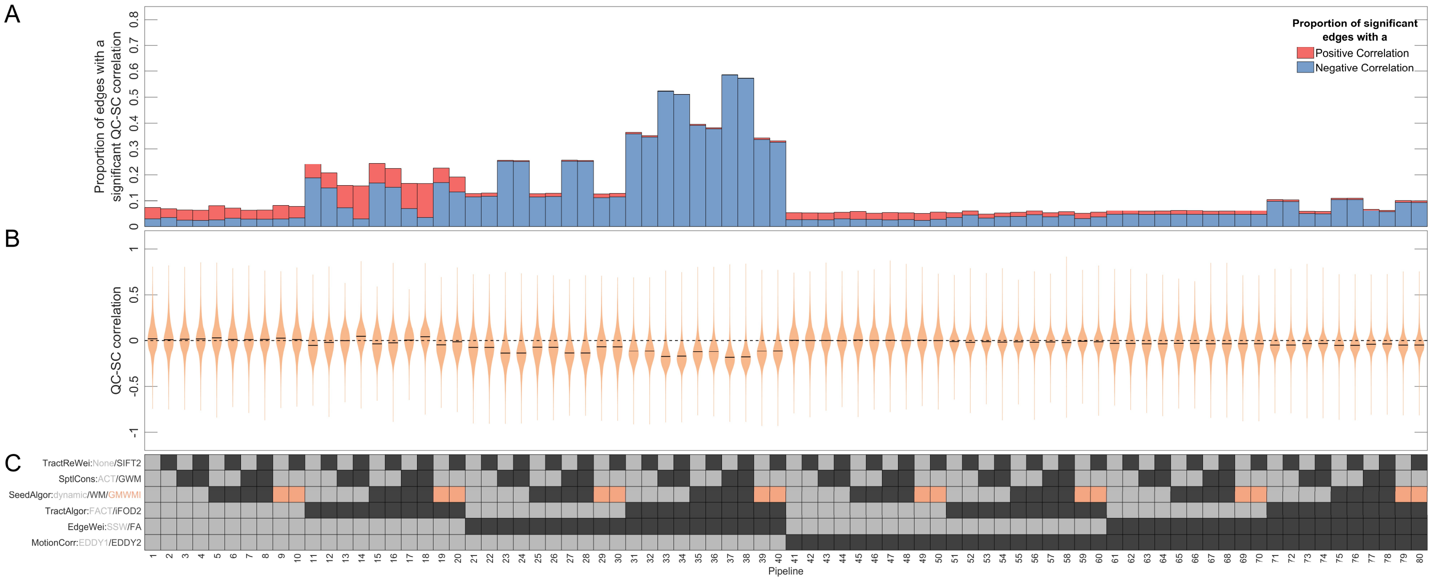


**Figure S5. The effect of in-scanner motion (*$ABS_{all}$*) on structural connectivity when using a 380 node parcellation.** (**A**) The proportion of edges (y-axis) that had a significant ($p$ < .05, uncorrected) QC-SC correlation in each pipeline (x-axis). Each bar is coloured to show out of those significant edges, what proportion were negative (blue) or positive (red). (**B**) The full distributions of QC-SC correlations (y-axis) for each pipeline (x-axis). (**C**) The preprocessing options used in each pipeline. Each row corresponds to a preprocessing step, with the possible options for that step colour-coded; the colour of the squares in each column indicate the specific methods used for a given pipeline.

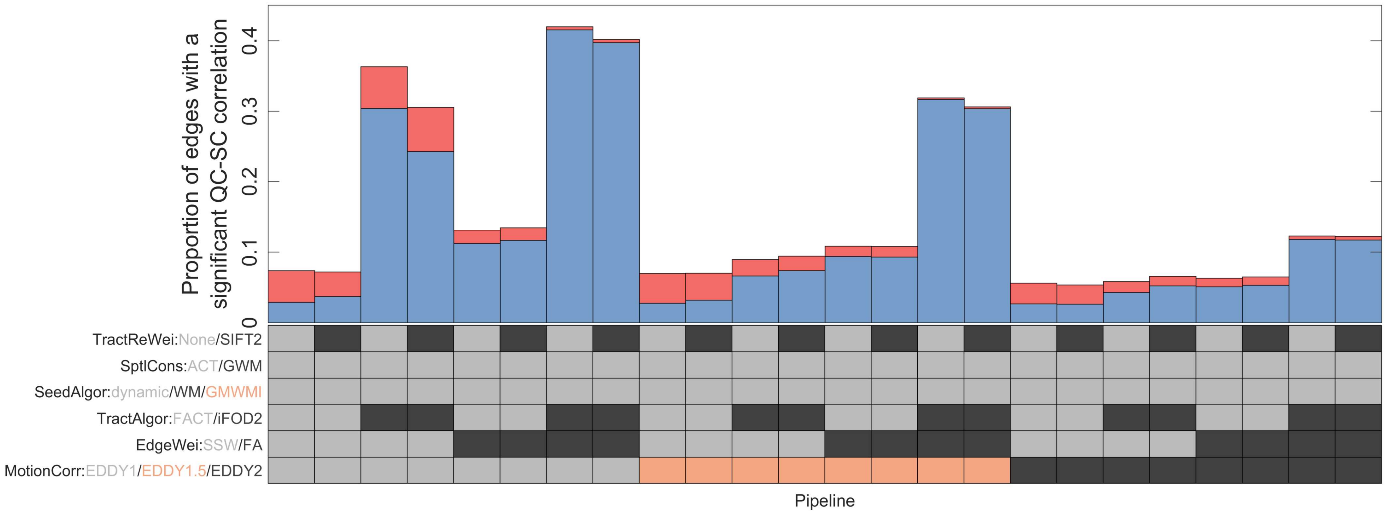


**Figure S6. The effect of in-scanner motion (*$ABS_{all}$*) on structural connectivity when using a 220 node parcellation, when using EDDY1, EDDY1.5, and EDDY2.** (**A**) The proportion of edges (y-axis) that had a significant (*p* < .05, uncorrected) QC-SC correlation in each pipeline (x-axis). Each bar is coloured to show out of those significant edges, what proportion were negative (blue) or positive (red). (**B**) The full distributions of QC-SC correlations (y-axis) for each pipeline (x-axis). (**C**) The preprocessing options used in each pipeline. Each row corresponds to a preprocessing step, with the possible options for that step colour-coded; the colour of the squares in each column indicate the specific methods used for a given pipeline. Note that pipelines using EDDY1.5 used a slightly different version of ACT (instead of a customised version it only used an image derived purely from FSL), but otherwise the pipelines were identical.

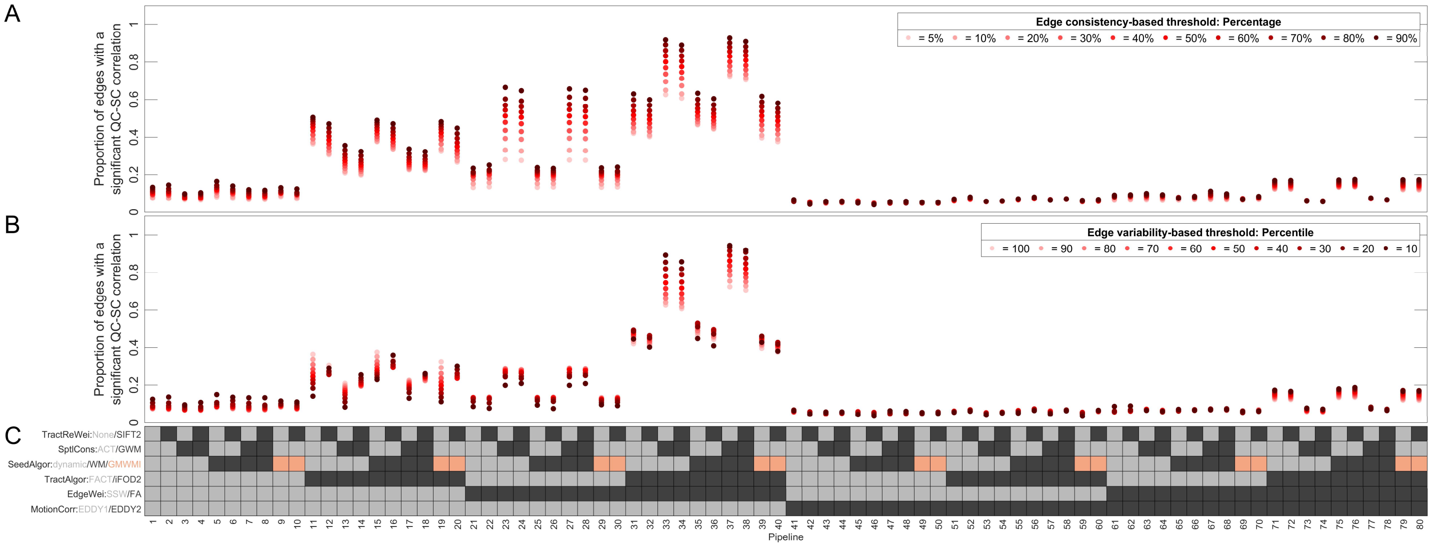


**Figure S7. The effect of in-scanner motion (*$ABS_{all}$*) on edge weight when using a 220 node parcellation in 80 combinations of processing choices across different edge consistency and variability thresholds.** (**A**) The percentage of edges (y-axis) which had a significant QC-SC correlation (y-axis) for each pipeline (x-axis) across edge consistency-based thresholds from 5% to 90%. (**B**) The percentage of edges (y-axis) which had a significant QC-SC correlation (y-axis) for each pipeline (x-axis) across edge variability-based thresholds from the 10$^{th}$ to 100$^{th}$ percentile. In (**A**) and (**B**) darker colours indicate a more stringent threshold. (**C**) The preprocessing options used in each pipeline. Each row corresponds to a preprocessing step, with the possible options for that step colour-coded; the colour of the squares in each column indicate the specific methods used for a given pipeline.

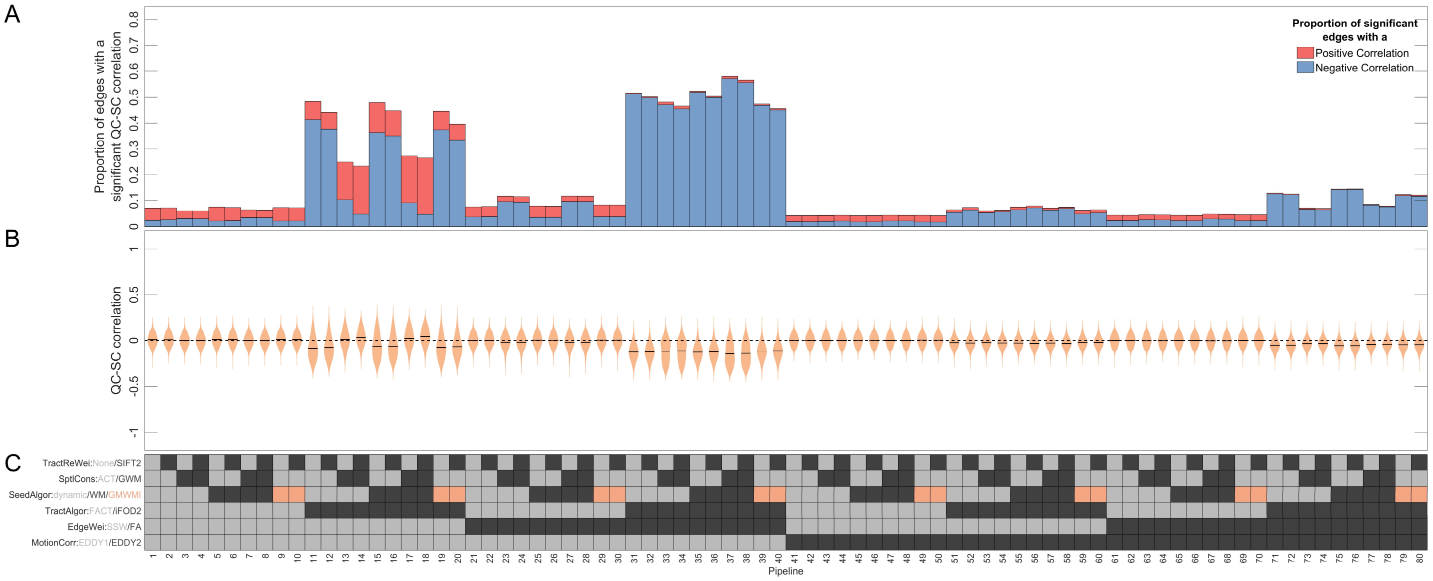


**Figure S8. The effect of in-scanner motion (*$ABS_{all}$*) on structural connectivity when using a 220 node parcellation, when not excluding subjects who had no connection.** (**A**) The proportion of edges (y-axis) that had a significant (*p* < .05, uncorrected) QC-SC correlation in each pipeline (x-axis). Each bar is coloured to show out of those significant edges, what proportion were negative (blue) or positive (red). (**B**) The full distributions of QC-SC correlations (y-axis) for each pipeline (x-axis). (**C**) The preprocessing options used in each pipeline. Each row corresponds to a preprocessing step, with the possible options for that step colour-coded; the colour of the squares in each column indicate the specific methods used for a given pipeline.

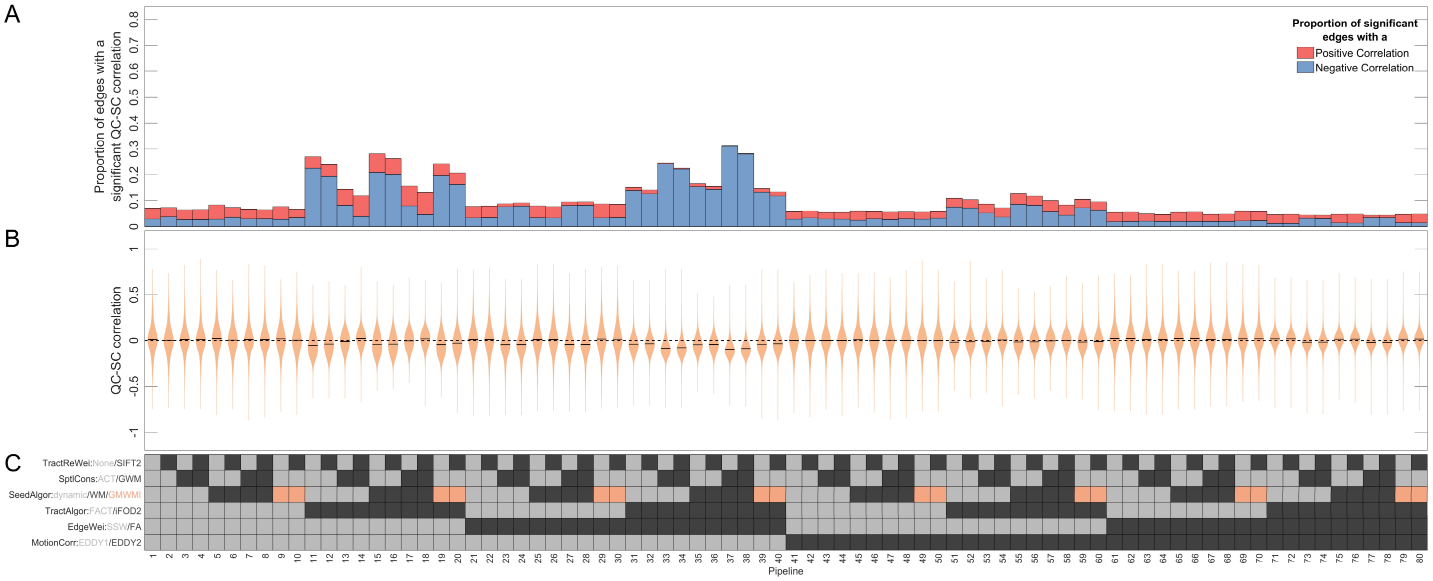


**Figure S9. The effect of in-scanner motion ($REL_{all}$) on structural connectivity when using a 220 node parcellation.** (**A**) The proportion of edges (y-axis) that had a significant ($p$ < .05, uncorrected) QC-SC correlation in each pipeline (x-axis). Each bar is coloured to show out of those significant edges, what proportion were negative (blue) or positive (red). (**B**) The full distributions of QC-SC correlations (y-axis) for each pipeline (x-axis). (**C**) The preprocessing options used in each pipeline. Each row corresponds to a preprocessing step, with the possible options for that step colour-coded; the colour of the squares in each column indicate the specific methods used for a given pipeline.

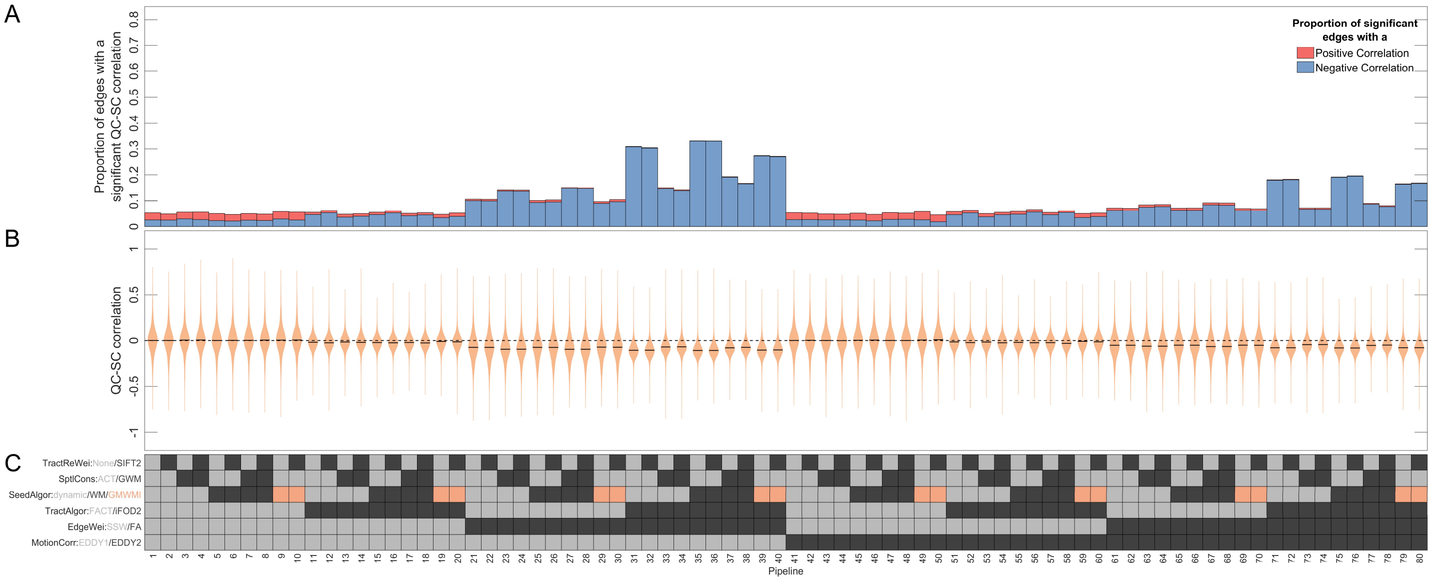


**Figure S10. The effect of in-scanner motion ($ABS_{b0}$) on structural connectivity when using a 220 node parcellation.** (**A**) The proportion of edges (y-axis) that had a significant ($p$ < .05, uncorrected) QC-SC correlation in each pipeline (x-axis). Each bar is coloured to show out of those significant edges, what proportion were negative (blue) or positive (red). (**B**) The full distributions of QC-SC correlations (y-axis) for each pipeline (x-axis). (**C**) The preprocessing options used in each pipeline. Each row corresponds to a preprocessing step, with the possible options for that step colour-coded; the colour of the squares in each column indicate the specific methods used for a given pipeline.

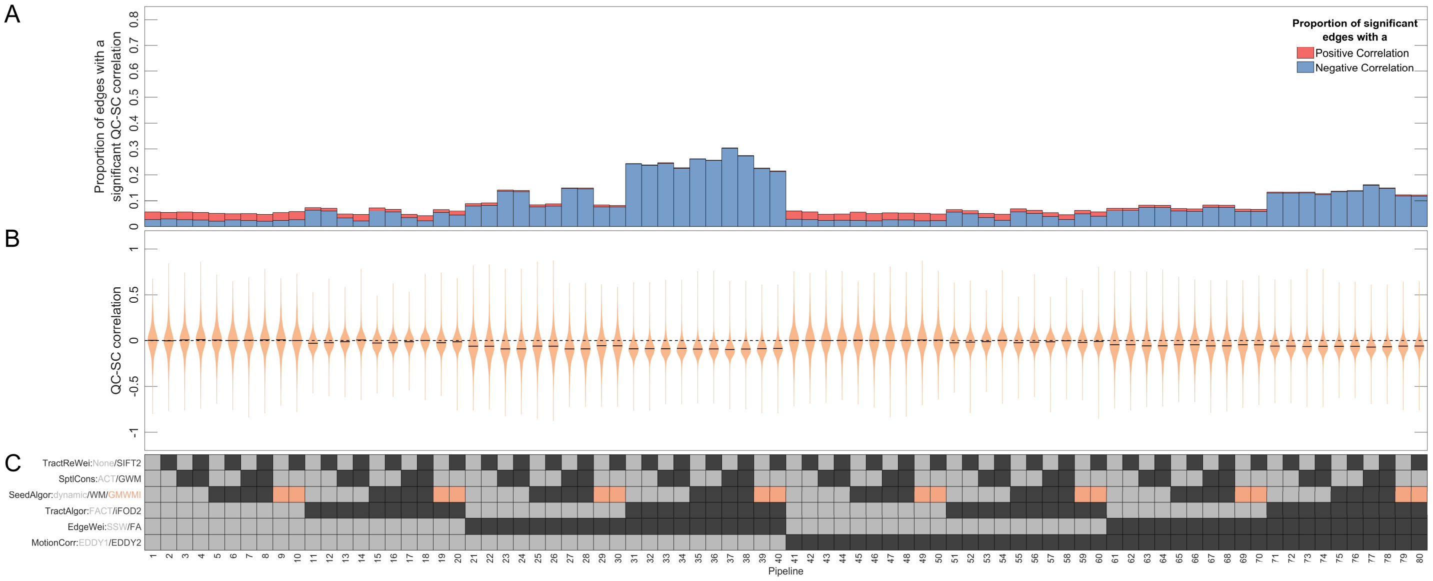


**Figure S11. The effect of in-scanner motion ($ABS_{b000}$) on structural connectivity when using a 220 node parcellation.** (**A**) The proportion of edges (y-axis) that had a significant ($p$ < .05, uncorrected) QC-SC correlation in each pipeline (x-axis). Each bar is coloured to show out of those significant edges, what proportion were negative (blue) or positive (red). (**B**) The full distributions of QC-SC correlations (y-axis) for each pipeline (x-axis). (**C**) The preprocessing options used in each pipeline. Each row corresponds to a preprocessing step, with the possible options for that step colour-coded; the colour of the squares in each column indicate the specific methods used for a given pipeline.

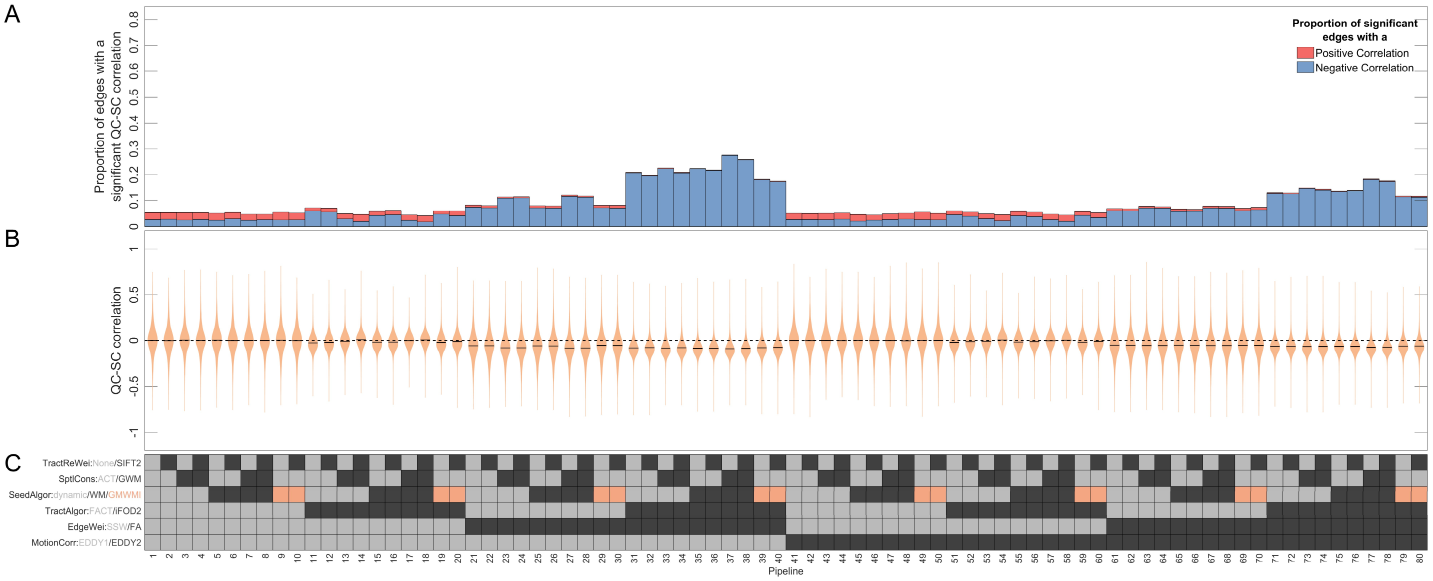


**Figure S12. The effect of in-scanner motion (*$REL_{b0}$*) on structural connectivity when using a 220 node parcellation.** (**A**) The proportion of edges (y-axis) that had a significant ($p$ < .05, uncorrected) QC-SC correlation in each pipeline (x-axis). Each bar is coloured to show out of those significant edges, what proportion were negative (blue) or positive (red). (**B**) The full distributions of QC-SC correlations (y-axis) for each pipeline (x-axis). (**C**) The preprocessing options used in each pipeline. Each row corresponds to a preprocessing step, with the possible options for that step colour-coded; the colour of the squares in each column indicate the specific methods used for a given pipeline.

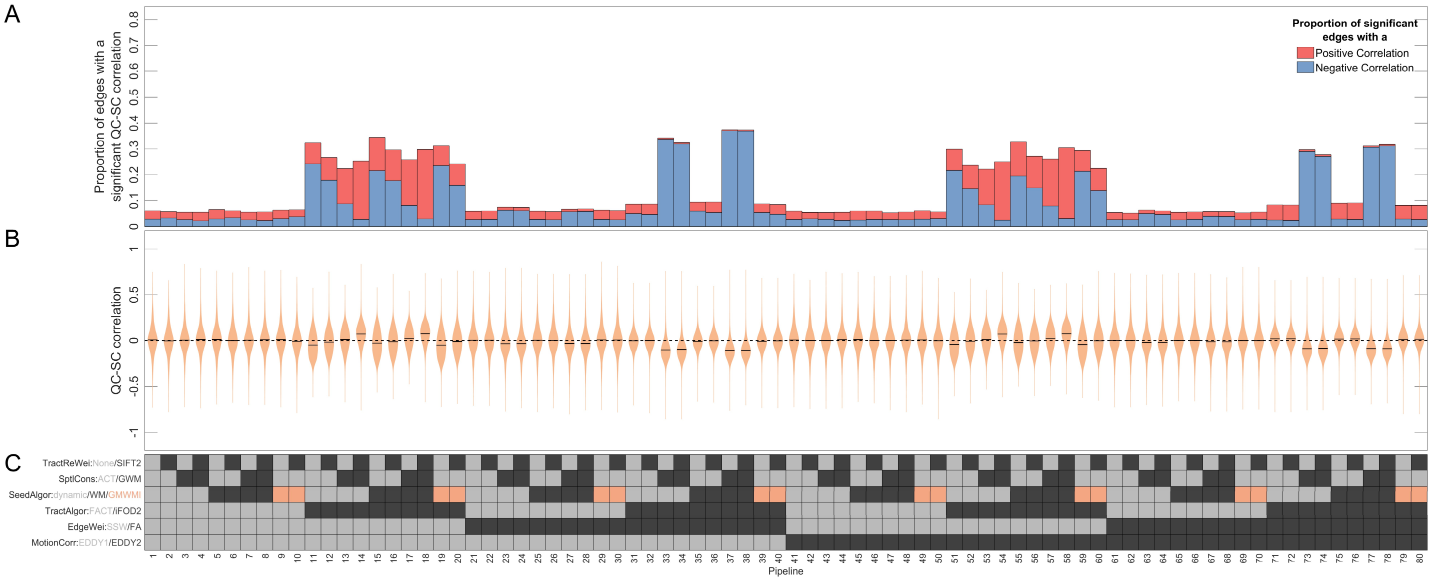


**Figure S13. The effect of in-scanner motion (*$REL_{b3000}$*) on structural connectivity when using a 220 node parcellation.** (**A**) The proportion of edges (y-axis) that had a significant ($p < .05$, uncorrected) QC-SC correlation in each pipeline (x-axis). Each bar is coloured to show out of those significant edges, what proportion were negative (blue) or positive (red). (**B**) The full distributions of QC-SC correlations (y-axis) for each pipeline (x-axis). (**C**) The preprocessing options used in each pipeline. Each row corresponds to a preprocessing step, with the possible options for that step colour-coded; the colour of the squares in each column indicate the specific methods used for a given pipeline.

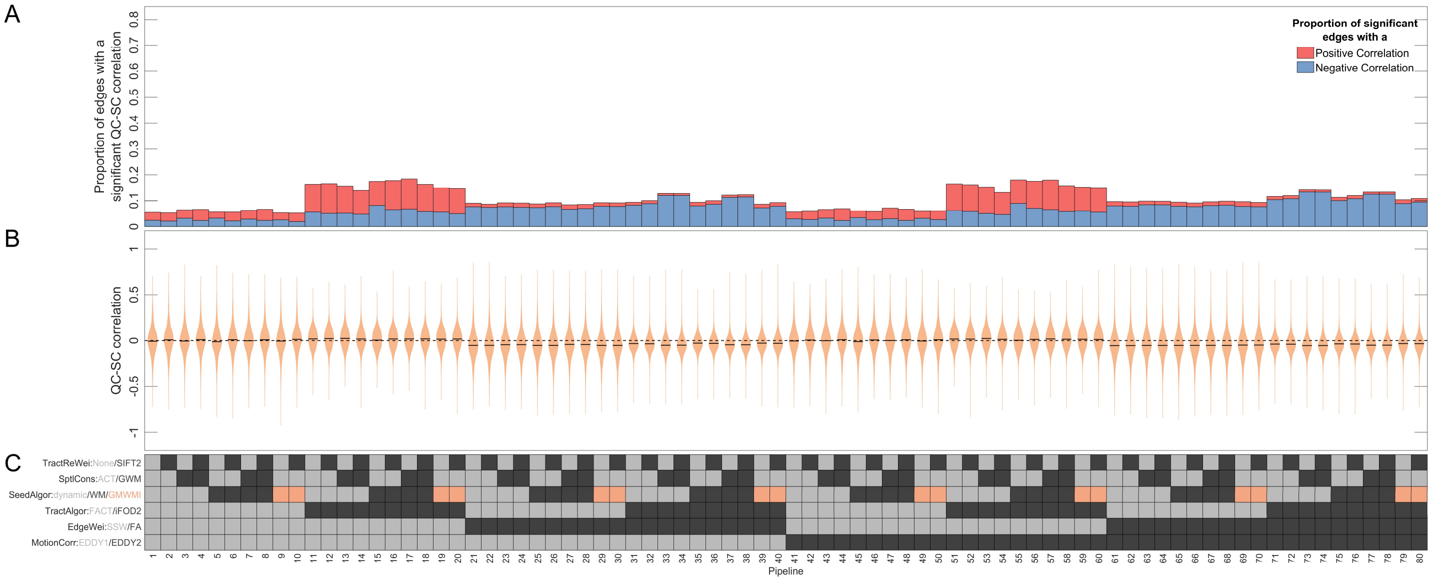


**Figure S14. The effect of in-scanner motion (*TSNR*) on structural connectivity when using a 220 node parcellation.** (**A**) The proportion of edges (y-axis) that had a significant ($p < .05$, uncorrected) QC-SC correlation in each pipeline (x-axis). Each bar is coloured to show out of those significant edges, what proportion were negative (blue) or positive (red). (**B**) The full distributions of QC-SC correlations (y-axis) for each pipeline (x-axis). (**C**) The preprocessing options used in each pipeline. Each row corresponds to a preprocessing step, with the possible options for that step colour-coded; the colour of the squares in each column indicate the specific methods used for a given pipeline.

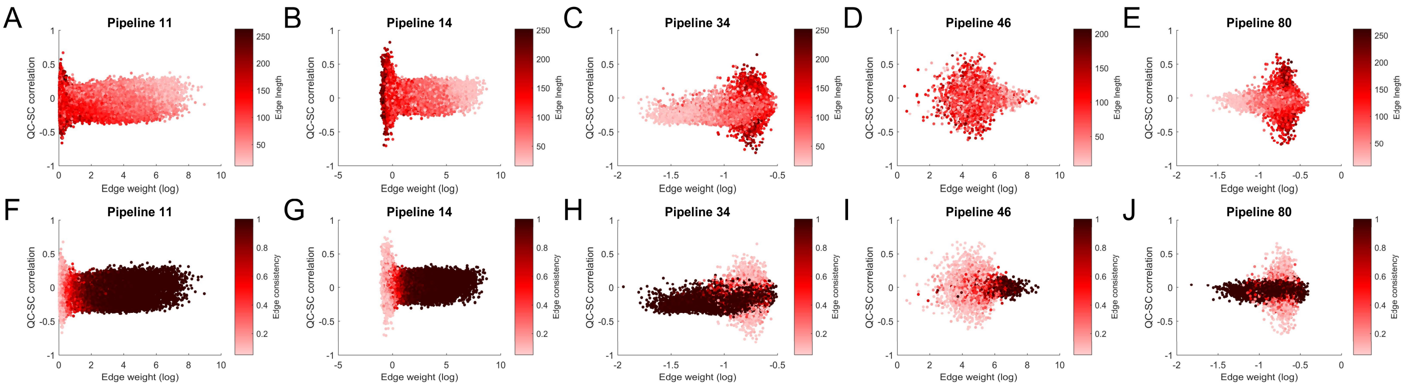


**Figure S15. Relationship of QC-SC correlations (*$ABS_{all}$*) and edge length, edge consistency, and edge weight.** The first row shows the relationship between QC-SC correlations (y-axis), edge weight (x-axis), and edge length (colourmap), while the second shows the relationship between QC-SC correlations (x-axis), edge weight (y-axis), and edge consistency (colourmap). Pipeline numbers correspond to those in Figures 2, 3, 5, and 7.

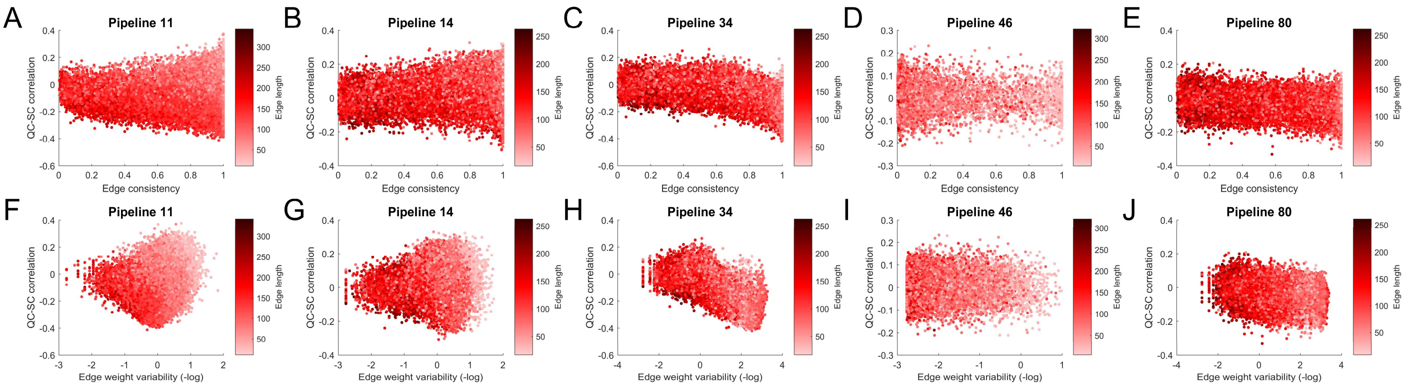


**Figure S16. Relationship of QC-SC correlations ($ABS_{all}$) and edge length, edge consistency, and edge weight variability when not excluding subjects who had no connection.** The first row shows the relationship between QC-SC correlations (y-axis), edge consistency (x-axis), and edge length (colourmap), while the second shows the relationship between QC-SC correlations (x-axis), edge weight variability (y-axis), and edge length (colourmap). Pipeline numbers correspond to those in Figures 2, 3, 5, and 7.

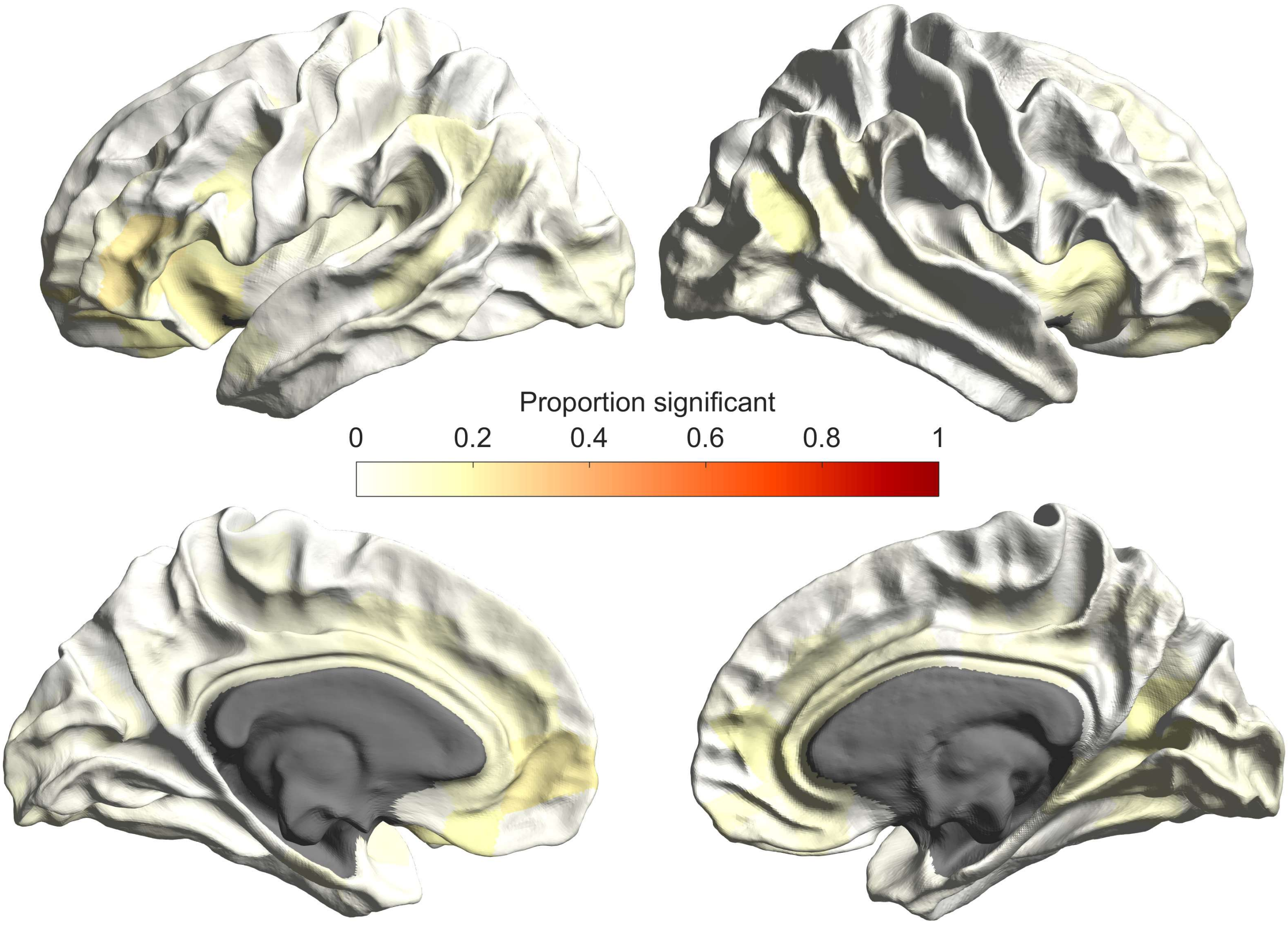


**Figure S17. Proportion of times across the 80 pipeline choices for the 220 node parcellations each node registered a significant QC-strength correlation.**

**Table S1.**

**Description of different motion measures**

| Measure | Abbreviation | Description |
|---|---|---|
| **Mean absolute volume-to-volume displacement** | $ABS_{all}$ | Mean root-mean-square of voxelwise displacements relative to the previous volume |
| **Mean relative volume-to-volume displacement** | $REL_{all}$ | Mean root-mean-square of voxelwise displacements relative to the first volume |
| **Mean absolute b = 0 volume-to-volume displacement** | $ABS_{b0}$ | Mean translation and rotation of b = 0 volumes across all three axes relative to the first b = 0 image |
| **Mean absolute b = 3000 volume-to-volume displacement** | $ABS_{3000}$ | Mean translation and rotation of b = 3000 volumes across all three axes relative to the first b = 3000 image |
| **Mean relative b = 0 volume-to-volume displacement** | $REL_{b0}$ | Mean translation and rotation of b = 0 volumes across all three axes relative to the previous b = 0 image |
| **Mean relative b = 3000 volume-to-volume displacement** | $REL_{b3000}$ | Mean translation and rotation of b = 3000 volumes across all three axes relative to the previous b = 3000 image |
| **Mean temporal signal-to-noise ratio of b = 3000 volumes** | *TSNR* | The mean signal-to-noise ratio as calculated across the b = 3000 volumes |

**Table S2.**

**Means and standard deviations of motion measures**

| Measure | M | SD |
|---|---|---|
| $ABS_{all}$ (EDDY1) | 1.05, uncorrected | 0.39 |
| $ABS_{all}$ (EDDY2) | 0.90 | 0.39 |
| $REL_{all}$ (EDDY1) | 0.61 | 0.32 |
| $REL_{all}$ (EDDY2) | 0.69 | 0.42 |
| $ABS_{b0}$ | 0.55 | 0.29 |
| $ABS_{3000}$ | 0.69 | 0.25 |
| $REL_{b0}$ | 0.32 | 0.14 |
| $REL_{b3000}$ | 0.58 | 0.13 |
| *TSNR* | 3.35 | 0.15 |
